# Evolution of the rate, molecular spectrum, and fitness effects of mutation under minimal selection in *Caenorhabditis elegans*

**DOI:** 10.1101/2025.09.24.678306

**Authors:** Sayran Saber, Md. Monjurul Islam Rifat, Fahimeh Rahimi, Michael Snyder, Amber Singh, Benjamin Eickwort, Yell Newhall, Moein Rajaei, Ayush Shekhar Saxena, Robyn E. Tanny, Vaishali Katju, Erik C. Andersen, Juannan Zhou, Charles F. Baer

## Abstract

The rate, molecular spectrum, and fitness effects of mutations vary at all levels of the biological hierarchy, from within individual genomes to among taxonomic domains. Understanding the evolutionary factors underpinning that variation is of fundamental importance to biology. Accurate quantification of the properties of mutations requires that other evolutionary forces, especially natural selection, be minimized as much as possible.

To investigate the evolution of the mutational process in *C. elegans*, we propagated a set of 100 “first order” mutation accumulation (O1MA) lines under minimal selection for ∼150 generations, divided each O1MA line into two “second order” MA (O2MA) lines and propagated them for another ∼150 generations, at which time the genome of each O2MA line was sequenced, and a subset of 50 O1MA families was assayed for competitive fitness.

Over the course of the experiment, the mean nucleotide substitution mutation rate did not change, but the variance increased. In contrast, the indel mutation rate increased significantly. The two types of mutations fulfill the predictions of different theoretical models for the evolution of mutation rate. These results reinforce previous findings that the rate of indels is more sensitive to endogenous stress than the rate of nucleotide substitutions.

Several evolutionary quandaries could be resolved if deleterious mutations interact synergistically (negative epistasis). Evidence for synergistic epistasis is famously inconclusive, although there is reason to think it may be more detectable under competitive conditions. However, a model of constant mutational effects on competitive fitness explains the results significantly better than a model including epistasis.

---

Evolution begins with mutation and would soon end if it stopped. Understanding the mutational process is thus of fundamental importance to all areas of evolutionary biology. Two distinct, but intertwined, lines of inquiry are (1): what factors influence the rate and molecular spectrum of mutation? and (2) what factors influence the effects of mutations on the phenotype and on fitness? In any case, to make progress it is necessary to consider a sufficiently large set of mutations unbiased by the effects of other evolutionary forces, natural selection in particular. In principle, this can be done either by investigating the properties of extremely rare segregating variants (Messer 2009; Schraiber et al. 2025), which most likely arose recently as new mutations and have been minimally scrutinized by natural selection, or by means of a laboratory mutation accumulation (MA) experiment, in which populations are manipulated in such a way as to minimize the efficacy of natural selection (Mukai 1964; Katju and Bergthorsson 2019). Each method has strengths and drawbacks. It is possible to characterize the sequence features of vastly more segregating variants than could ever be accumulated in an MA experiment, and those mutations integrate over genotype and environment in (recent) time and space. However, the phenotypic effects of very rare segregating variants are usually not measurable, and the fitness effects are estimatable only indirectly and are confounded with the effects of demography (Johri et al. 2022).

MA experiments, on the other hand, have the significant limitation that only a comparatively small number of mutations can be accumulated on at most a few genetic backgrounds in at most a few environmental contexts that are assuredly unnatural for the organism. However, MA experiments come with two major advantages: first, that the experimenter controls demography (and thus the efficacy of selection) and the environment, and second, that the effects on phenotype and fitness can be measured directly, albeit in the lab environment and (usually) only in aggregate.

It is by now well-established that the rate and spectrum of mutation vary even within species and populations (Wang and Obbard 2023; Lynch et al. 2023; Sasani et al. 2022; Bergeron et al. 2023), and the causes of that variation are beginning to be disentangled (Sasani et al. 2024; Wei et al. 2022; Liu and Zhang 2019). Most studies designed to parse the causes of variation in the mutational process have been done in microbes, primarily for reasons of tractability, although relevant studies have been done in a variety of multicellular organisms, including humans (Carlson et al. 2018; Carlson et al. 2020; Garcia-Salinas et al. 2025). A few broad-scale patterns have emerged, among them (*i*) the per-genome, per-generation mutation rate (*U*) varies inversely with population size and body size, at least at large phylogenetic scale (Lynch et al. 2023), (*ii*) the per-nucleotide, per-generation mutation rate (μ) varies inversely with genome size in microbes (“Drake’s Rule”, Drake 1991) but positively with genome size in multicellular eukaryotes (Lynch et al. 2016), and (*iii*) the mutation rate often increases under conditions of physiological stress (Williams and Foster 2012; Ram and Hadany 2012; Liu and Zhang 2019). Obviously, however, much remains to be learned about the causes of variability in the mutational process.

A second feature of the mutational process of longstanding interest is the distribution of fitness effects (DFE) of new mutations (Fisher 1930; Charlesworth 1996; Eyre-Walker and Keightley 2007). One feature of the DFE is certain, and another nearly so. First, it is beyond doubt that most mutations with non-trivial effects on fitness are deleterious. Second, in diploid organisms, there is abundant evidence for a negative relationship between the strength of selection and the degree of dominance: on average, the more deleterious the mutation, the closer it is to being completely recessive (Simmons and Crow 1977; Di and Lohmueller 2024). Beyond that, the picture gets murky.

For several reasons, it is of great interest to know how mutations, especially deleterious mutations, interact *on average*. Whether the average interaction between deleterious mutations is multiplicative (no epistasis), synergistic (negative epistasis) or diminishing-returns (positive epistasis) has important implications for evolution, prominently including the evolution of sex and recombination (Barton 1995; Otto 2009), as well as the long-term existence of species with modest effective population sizes (e.g., humans, Kondrashov 1995; Charlesworth 2013b). The evident long-term advantage of sex and recombination becomes easier to explain if deleterious mutations interact synergistically, as does our continued existence.

For related reasons, it is also of considerable interest to understand the relationship between environmental stress and the fitness effects of mutations. There is a longstanding general intuition that mutations become more deleterious under stress, but as trenchantly pointed out by Agrawal and Whitlock (2010), there is no a priori theoretical justification for that intuition, although they convincingly argue that deleterious effects are likely to be magnified under competitive conditions.

To experimentally investigate the evolution of the mutational process in the (near) absence of natural selection, we employ what we refer to as a “second-order” MA experiment (Figure 1A). The O2MA design lets us quantify the evolution of the rate and spectrum of mutation, as well as the evolution of mutational effects on fitness. The resulting data allow us to address several fundamental questions in evolutionary biology. Of specific interest are (*i*) the cumulative directional effect of new (anti)mutator alleles (i.e., the mutational bias for mutation rate), and (*ii*) the rate of input of new genetic variance for mutation rate (mutators and antimutators). The relative frequency of (anti)mutators has important implications for the evolution of mutation rate (Drake 1993; Lynch 2011), and it is commonly assumed that mutator alleles arise by spontaneous mutation more often than antimutators (e.g., Lynch et al. 2016).

**Figure 1.**
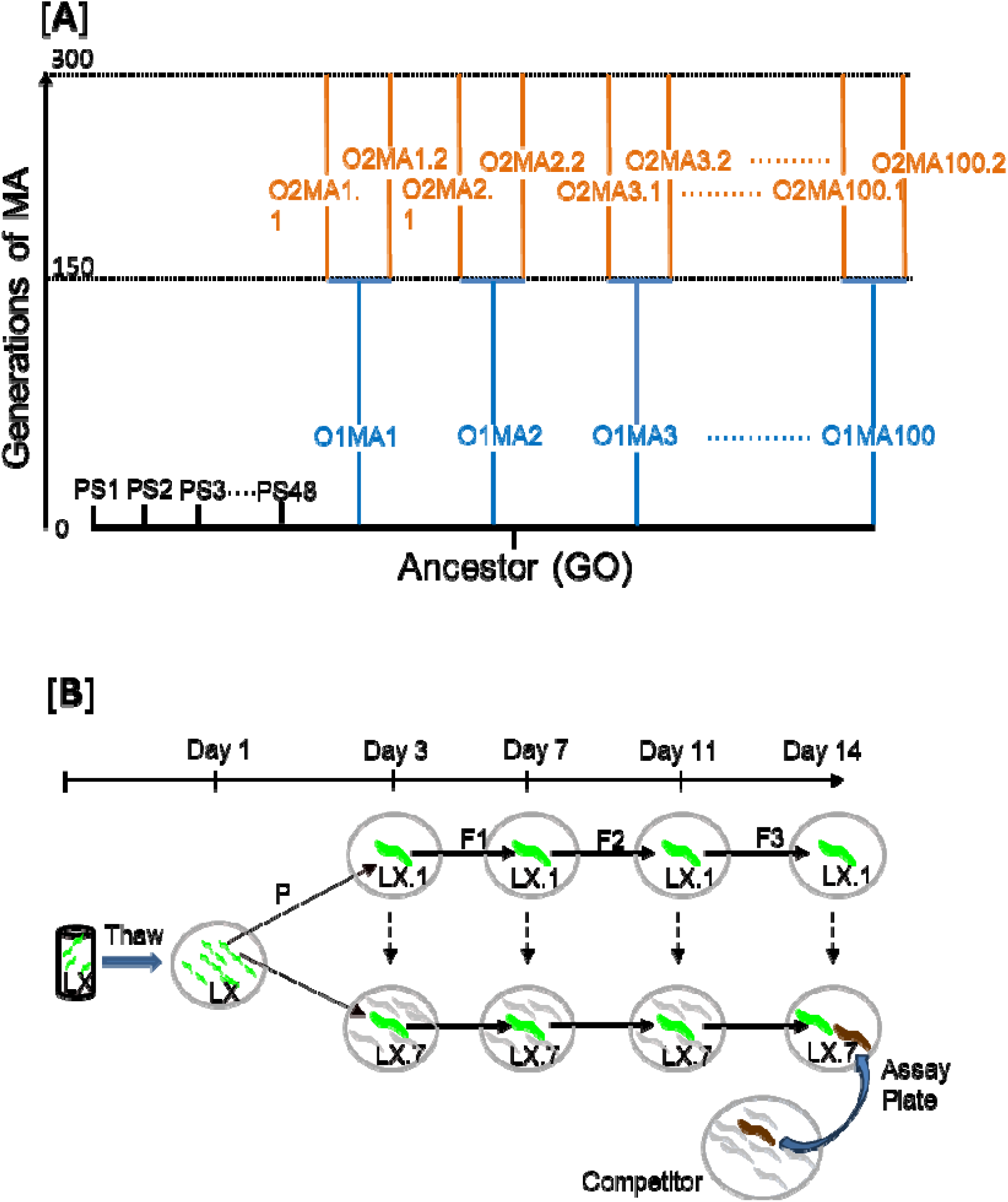
(**A**) Schematic depiction of the MA experiment. Short black lines depict ancestral G0 “pseudolines” (PS), blue lines depict O1MA lines, orange lines depict O2MA lines. (**B**) Schematic depiction of the competitive fitness assay. Label “LX” refers to “Line X” (MA or G0), label LX.1,…LX.7 refers to replicates 1-7 of a line in the assay. See Methods for details.

However, there is little quantitative information with respect to the rate and bias of spontaneous mutations affecting mutation rate, especially in multicellular organisms, and a recent study in yeast reported a surprisingly high frequency of spontaneous antimutators (Liu and Zhang 2021).

Also of interest is (*iii*) the relationship between mutation rate and fitness. If the optimum mutation rate is zero (Sturtevant 1937; Kimura 1967; Lynch 2008), the expectation is that fitness will always be negatively correlated with mutation rate. However, if the mutation rate is under stabilizing selection with an optimum greater than zero (Kimura 1967; Fitzsimmons et al. 2018), the implication is that genotypes with intermediate mutation rates will have higher fitness than genotypes with more extreme mutation rates. Further, if the mutation rate is condition-dependent, the mutation rate may itself be an increasing function of the mutation load, which has implications for both the evolution of sex and recombination (Agrawal 2002) and the survival of small populations (Shaw and Baer 2011).

Finally, we are interested in (*iv*) the average interaction between the accumulated mutations with respect to fitness. The O2MA design allows us to quantify the average mutational effect as a function of the mutation load, and thus the average direction of epistasis. The average epistatic effect of new mutations has been of interest to population geneticists for over fifty years (Mukai 1969), but the empirical evidence is frustratingly inconclusive (de Visser et al. 2011). However, there is some reason to believe that epistasis is magnified under competitive conditions (Agrawal and Whitlock 2012; Peck and Waxman 2000), and most empirical investigations of mutational epistasis have not measured competitive fitness.

Items (*i*) and (*ii*) are addressed by means of whole-genome sequencing; items (*iii*) and (*iv*) are addressed using experimental assays of competitive fitness. We elaborate on each of these topics in the Discussion.

## Methods and Materials

### Mutation Accumulation (MA) Experiment

We obtained a stock of the EG8072 strain of *C. elegans* (https://cgc.umn.edu/strain/EG8072) from the Caenorhabditis Genetics Center (CGC) at the University of Minnesota in September 2018. EG8072 is a derivative of the standard N2 strain, with green (GFP) and red (mCherry) fluorescent reporters stably integrated into the genome. We propagated the founder stock for seven generations by transfer of a single immature hermaphrodite at 4-day intervals, at which point the population is expected to have reached mutation-drift equilibrium (6*N_e_*generations; Lynch and Hill 1986). The population was expanded to large size (thousands of mixed-stage individuals) for 2-3 generations and cryopreserved.

In November 2018 we thawed an aliquot of the cryopreserved stock, cleaned it by bleaching (Stiernagle 2006) and initiated 148 replicate populations, each initiated from a single immature hermaphrodite. 48 replicates were expanded to large size (N ≈ a few thousand) for an additional 2-3 generations and cryopreserved in 20 replicate PCR plates. These are the G0 ancestral “pseudolines”. The remaining 100 populations – the first-order (O1) MA lines - were propagated by transfer of a single immature hermaphrodite at four-day intervals for an additional 600 days (Gmax=150 generations). MA lines were maintained under standard conditions: on 35 mm NGMA agar plates spotted with 50 µl of an overnight culture of *E. coli* OP50, kept in a Percival incubator set to a constant 20° C. Plates were kept on cafeteria trays, in 6 X 8 arrays, 48 plates per tray. Even numbered and odd numbered replicates were kept on different trays.

After the leading-generation worm was transferred, the plate it was taken from was moved to a different incubator set to 10° C; this is the (first) backup plate. Worms do not reproduce (or reproduce only very slowly) at 10° C. If at the time of transfer a worm of the leading generation failed to reproduce or was not visibly gravid, it was replaced by a sibling taken from the previous generation plate. If there were offspring on the leading generation plate but no worm had reached at least the L2 stage, the plate was “held over” without transferring a worm. This protocol assures that generation number of a line is known with (near) certainty. Demographic data on backups and holdovers is given in Supplementary Table 1.

All lines were cryopreserved at 50 generation intervals. After 150 transfers (600 days, Gmax=150), each O1MA line was replicated into two second-order (O2) MA lines; the two O2MA lines derived from an O1MA line are an “O1MA family”. The O2MA lines were propagated as described for an additional ∼150 generations (Gmax=306), at which time they were cryopreserved analogously to the G0 pseudolines as described above.

### Genome Sequencing and Variant Calling

#### (i) DNA Extraction and library preparation

Cryopreserved tubes of worms were thawed and grown for several days on 60 mm NGMA agar plates containing Streptomycin and Nystatin and seeded with 110 μl of overnight culture of the HB101 strain of *E. coli*. When food was nearly exhausted, a small piece (∼2 cm^2^) of the agar plate was transferred onto a 100 mm NGMA plate and worms were allowed to grow for another 2-3 days. When food was nearly exhausted, worms were collected into 15 ml centrifuge tubes on ice and allowed to settle. Approximately 100 μl of settled worms were transferred into a 1.5 ml microcentrifuge tube and stored at -80°C until all samples were ready for DNA extraction, at which time samples were thawed, and genomic DNA was extracted with the DNeasy Kit (Qiagen) following the manufacturer’s protocol. Extracted DNA was stored at -20°C prior to library preparation.

Frozen DNA samples were thawed and diluted with diH_2_O to a concentration of 0.2 ng/μl. Tagmentation was performed per manufacturer’s instructions (Illumina, Nextera DNA Sample preparation kit, FC-121-1030). Following tagmentation, amplification was performed using custom barcoded IDT primers. Following amplification, a 96-pooled well sample was generated. Once pooled, 170µL of sample was combined with 30µL of 6X loading dye and run on a 2% agarose gel. DNA segments of 300-500 bp were excised from gel. The QIAquick Gel Extraction Kit (QIAquick, 28704) was used to clean and elute DNA from the gel, following manufacturer’s instructions. Pooled library samples were quantified using the Qubit HS kit (Qubit, Q33230) and sequenced to ∼25X average depth of coverage with 150 bp paired-end Illumina sequencing on a NovaSeq 6000 by Novogene, Inc.

#### (ii) Genome Assembly

Adapter sequences were trimmed from the raw sequence reads with fastp (Chen et al. 2018). Reads were then aligned to the *C. elegans* N2 strain reference genome (WS292) using bowtie2 (Langmead and Salzberg 2012). The resulting BAM files were sorted and filtered with Samtools (Li et al. 2009) and bamTools (Barnett et al. 2011) for downstream processing. Read group information was added with Picard (http://broadinstitute.github.io/picard). Duplicate reads were identified and removed using MarkDuplicates in GATK (McKenna et al. 2010). Sequence quality was assessed with FastQC (Andrews 2010) and MultiQC (Ewels et al. 2016) after trimming and alignment and prior to variant calling.

#### (iii) Variant Calling Pipeline

Variants were called with the HaplotypeCaller tool in GATK (McKenna et al. 2010) run in GVCF mode. HaplotypeCaller identifies both single nucleotide variants (SNVs) and indels by local de novo assembly of haplotypes in regions with evidence of variation, improving accuracy by reconstructing the sequence context in those regions. GVCFs from all samples were merged with GenomicsDBImport, and joint genotyping was performed using GenotypeGVCFs. Sites were filtered using GATK VariantFiltration to retain genotypes with a minimum coverage depth of either DP ≥ 3 or DP ≥ 10. These two filtered datasets were then analyzed in parallel in all subsequent steps. We used bcftools (Li 2011) to identify homozygous variants unique to a line, which were counted as putative mutations.

Variants that met the following three criteria were considered as putative mutations: (1) they were called homozygous; (2) they were present in one and only one O1MA family, and (3) the ancestor (G0) genotype is homozygous wild-type. If the variant is present in both O2MA sublines, the mutation is inferred to have occurred in the first 150 generations and be a first-order (O1) mutation. If the variant is present in only one of the two O2MA sublines, it is inferred to have occurred subsequent to the split of the two O2MA sublines and be a second-order (O2) mutation. Comparisons of O1 and O2 mutation rates were restricted with the additional criteria (4) a site must be called in both O2MA lines in an O1MA family, (5) the site must be called in at least 90% of O1MA families, and (6) variants within 100 bp of another variant (“multi-nucleotide variants”, MNV) in the same O1MA family were omitted from the analysis of SNVs and indels. MNVs are analyzed as a separate category of mutation.

One O2MA line from O1MA family 9 was not sequenced. The identities of two O2MA lines appear to have been swapped, one from O1MA family 34 and one from family 46. One O1MA line (family 26) evolved an extreme indel mutator phenotype, as evidenced by the >5X greater indel rates of the two O2MA sublines than of the O2MA line with the next highest indel rate (Supplementary Table 2). Investigation of the mutations in family 26 revealed two potential O1 mutator mutations in the genes *chd-3* and *mut-16*. *chd-3* is a chromodomain helicase binding protein involved in the repair of meiotic double-strand breaks (Turcotte et al. 2018); *mut-16* is involved in (among other things) nuclear siRNA suppression of TEs (Zhang et al. 2011). Except where noted, those families were omitted from further analysis.

In the O2MA lines, over 97% of the ∼100.2 Mb genome was covered at the liberal 3X coverage criterion (mean= 97.6%, median = 97.8%). At 3X coverage, the mean and median fractions of the genome included in the O1MA analysis are 97.3% and 97.5% respectively. At 10X coverage the O2MA mean and median are 93.5% and 95.8%; the O1MA mean and median are 89.8% and 93.2% (Supplementary Table 2).

### Assessment of falsely inferred variants

#### (i) False positives

EG8072 is derived from the N2 strain, which is the source of the *C. elegans* reference genome (Ichikawa et al. 2025). Any difference between our EG8072 ancestor and the reference genome is either (1) a mutation that fixed subsequent to the divergence of EG8072 and the N2 reference strain, (2) a false negative (FN) in the N2 reference, or (3) a false positive (FP) in our EG8072 ancestor. In the first two cases, all MA lines derived from the EG8072 ancestor should also have the variant allele. However, if the variant is FP in our EG8072 ancestor, the derived MA lines should all have the reference allele. The EG8072 G0 ancestor was sequenced from three independent Illumina libraries derived from the same extracted DNA sample. We estimated the frequency of FPs from each of the three samples of the ancestor.

At the liberal (3X) coverage criterion we identified an average of 1662 fixed variants between our G0 ancestor and the reference genome (1676, 1647, and 1664 in the three replicates, respectively; Supplementary Table 3), of which approximately 40% were single nucleotide variants (SNVs), 37% were insertions, and 22% were deletions. Approximately 20% of those variants are multi-nucleotide variants (MVNs), defined as any variant occurring within 100 bases of another variant. Proportions of variants in an MNV that are SNVs, insertions, and deletions are 53%, 28% and 20%, respectively. In the three G0 replicates we identified three (1 deletion, 2 SNVs), three (3 SNVs) and one (1 SNV) FPs respectively. One of the FP SNVs (a G:C→C:G transversion) was common to all three replicates; the others were only found in one replicate. The sole deletion is in an MNV within 10 bp of an SNV in the same replicate line. Of the seven FPs detected at the liberal 3X criterion, six were also found at the stringent 10X criterion. By way of comparison, the same analysis with a different N2 ancestor found one FP (an SNV) out of 1439 variants in one replicate of the G0 ancestor and 68 MA lines at the 3X criterion (Rajaei et al. 2021).

#### (ii) Failure to recall (FtR) and false negatives (FN)

A mutation can fail to be detected (“failure to recall”, FtR) for two reasons. First, a site with a true mutant may not be called (i.e., missing data). Second, a true mutant allele may be mistakenly identified as an ancestral allele; that is a “true” false negative (FN). FNs are a subset of FtR and cause the mutation rate to be underestimated. FtR due to missing data may or may not bias estimation of the mutation rate, depending on whether sites with true mutations are more or less likely to be called than sites without mutations. All else equal, sites with true mutations are less likely to be called than sites without mutations. The reason is that if a sequence with *n* or more mismatches exceeds the cutoff for mapping, a sequence with a mutation has one more mismatch than the same sequence without the mutation. All else also equal, FNs are equally likely to be true O1 mutations or true O2 mutations. Since in any sequenced O2MA line an FN is equally likely to be an O1 or O2 mutation (Figure 2) FNs do not introduce a systematic bias (assuming equal O1 and O2 mutation rates).

**Figure 2.**
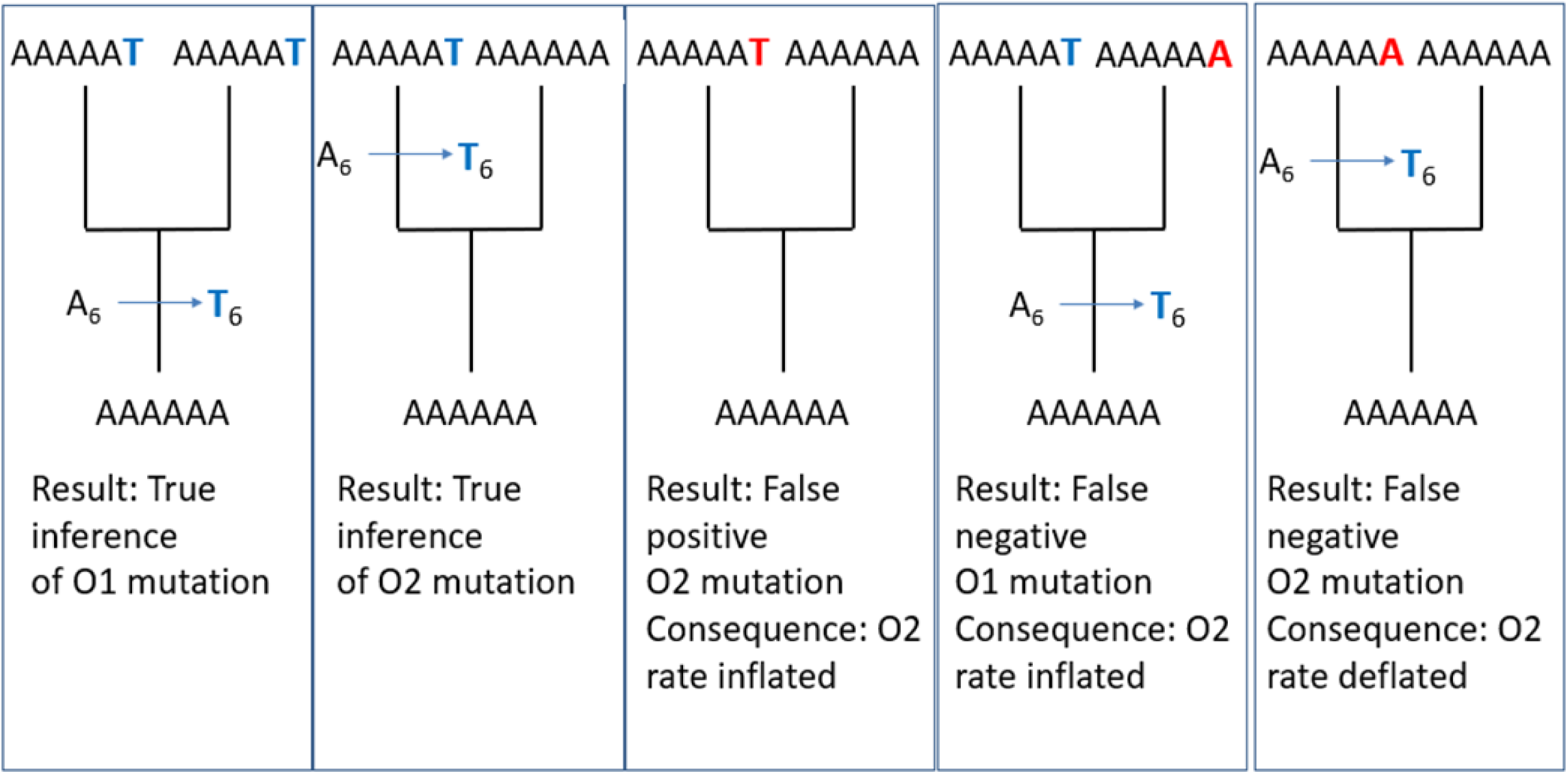
Schematic depiction of the effects of false inferences. Stem branch is the O1MA line, bifurcating terminal branches are the two O2MA lines derived from the O1MA line. Mutations are shown represented as arrows on the lineage. True sequence inferences are shown in blue text at sequence position 6; false inferences are shown in red text at sequence position 6.

To assess our ability to recall mutations, we introduced simulated “dummy” mutations at random into the reference genome and analyzed the data using our standard variant-calling pipeline, but with the simulated genome as the reference (Rajaei et al. 2021). The changed site in the reference should appear as a variant in our sequenced sample. For example, suppose the true reference allele at a position is an A and we change it to a C. The site should be called variant (A/A in this example) in our data. If the site is either not covered in the data or is called anything other than a variant homozygote (A/A), we have failed to recall the mutation. If the site is covered in the data but not called a variant homozygote, that is an FN.

We constructed a pseudo-reference genome by inserting 1200 single-nucleotide variants (SNVs) and 12,000 indels (1–20 bp, size chosen at random) into the *C. elegans* N2 (WS292) reference genome (Supplementary Table 4). First, we randomly inserted “dummy” SNVs and indels into the reference genome in random positions using the R packages GenomicRanges, IRange, and Biostrings and compiled them into a VCF file. The VCF was then sorted and indexed with samtools, and the pseudo-reference genome was produced with GATK FastaAlternateReferenceMaker. The genome assembly and variant calling pipelines were rerun with all the MA lines and the G0 ancestor using this pseudo-reference genome. Line-specific simulated FtR and FN rates and counts are given in Supplementary Table 5.

To validate the consistency of the procedure, the original variants (i.e., putative mutations) were re-mapped against the pseudo-reference genome. All sites identified as variant in the original analysis were also called as variant against the pseudo-reference. We repeated the simulation twice independently and obtained consistent results across runs (Supplementary Table 6).

At the liberal 3X criterion, the average FtR rate was ∼2.2% for SNVs and ∼2.5% for indels. The FN rates were 0.21% and 0.53% for SNVs and indels. At the stringent 10X criterion the failure to recall rates were 7.8% and 7.7% for SNVs and indels respectively; the FN rates were 0.084% and 0.12%. All analyses are based on the 3X coverage data; 10X data are reported in the Supplementary Tables.

### Competitive Fitness Assay

#### (i) Focal worms

50 G150 O1MA lines and 20 G0 pseudolines (PS lines) were chosen at random for inclusion in assay block 1. Block 2 included one G300 O2MA replicate from each of those same 50 O1MA families and the same 20 PS lines; Block 3 included the other O2MA replicate from each of the same 50 O1MA families, along with the same 20 PS lines. At the beginning of each assay (*Day 1*) a cryopreserved aliquot from each line (MA and PS) was thawed, and ∼150 µl pipetted onto a 60 mm NGMA plate seeded with the 100 µl of an overnight culture of the OP50 strain of *E. coli* (the thaw plate; Figure 1B) Two days later (*Day 3*), seven immature worms (L3/L4 stage) from each line were picked singly to a 35 mm NGMA plate seeded with 50 µl of OP50. These plates are the parental (P) generation. Each P plate was identified with a random number and arranged in numerical order on cafeteria trays, 48 plates/tray, and placed in a Percival incubator maintained at 20°C and ambient humidity. Thaw plates were incubated at 10°C to prevent further reproduction. Four days later (*Day 7*), a single immature (L3/L4) hermaphrodite was picked to a new plate and maintained as before. These plates are the F1 generation. If a P generation worm failed to reproduce, it was replaced in the assay with a sibling from the thaw plate. Four days later (*Day 11*) the process was repeated for the F2 generation. If an F1 worm failed to reproduce, the replicate was removed from the assay and treated as missing data. Three days later (*Day 14*) a single L1 stage worm was picked from each F2 plate onto the assay plate, along with an L1 competitor strain worm.

#### (ii) Competitor worms

On *Day 1* of the assay, an aliquot of N2 strain worms was thawed and pipetted onto a seeded 35 mm plate and allowed to reproduce. When the food was exhausted (around *Day 5*), a ∼2 cm chunk was transferred to a new 100 mm NGMA plate seeded with 400 µl of OP50 and allowed to reproduce. When food was exhausted (around *Day 9*), a chunk was transferred to a new plate. On *Day 13*, worms were washed from the plate and bleached to synchronize the population. Bleached embryos were pipetted in equal aliquots onto four 100 mm NGMA plates supplemented with streptomycin and the antifungal agent nystatin and seeded with 400 µl of the HB101 strain of *E. coli*.

#### (iii) Assay plates

A single L1 focal worm (MA or G0 pseudoline) and a single L1 competitor worm were picked to a 35 mm NGMA plate supplemented with streptomycin and nystatin and seeded with 50 µl of HB101. Assay plates were identified by random number, placed in random order on trays (48 plates/tray) and incubated at 20°C for eight days (*Day 21*; approximately two overlapping generations), at which point nearly all plates were depleted of bacterial food.

Worms were washed from the assay plate in 2 ml of M9 buffer and transferred to a 2 ml deep-well 96-well plate. The plate was centrifuged for 1 min at 1,000 g, and the supernatant was aspirated with an 8-channel strip aspirator set to leave 100 μl in each well. The worms were washed once more by adding 1.4 ml of M9 buffer, centrifuging, and aspirating as before, followed by the addition of 1.4 ml of M9. Worms were mixed by gently pipetting with a 12-channel pipette and a 110 μl sample was transferred into a well in a 96-well, clear-bottom tissue culture plate. 5 mM levamisole was added to each well to immobilize worms prior to imaging.

#### (iv) Imaging

Bright-field and GFP fluorescence images (470±20 nm excitation, 525±25 nm emission) were captured at 20x or 30x magnification for each well using an automated epifluorescence microscope (IX-70, Olympus, Pittsburgh, PA, USA) fitted with a CCD camera (Retiga-2000R, QImaging, Surrey, BC, Canada), XYZ stage and focus motors (Prior Scientific, Cambridge, UK), and controlled by Image Pro-plus software (Media Cybernetics, Rockville, MD, USA). 96-well plates were imaged first under bright field (BF) and then under fluorescence (GFP). Images of the same well (BF and GFP) were paired and renamed after checking by eye that the images matched.

#### (v) Counting

Worms were counted from stacked images using ImageJ software. We summarize the protocol here; a detailed tutorial of the ImageJ analysis is included as Supplementary Appendix 1.

*Step 1*. Paired images were stacked using the “Stacks” option under the “Image” tab.

*Step 2*. Images were merged and colorized using the “Merge Channels” tool under the “Image” tab, with fluorescent worms colored differently from non-fluorescent ones.

*Step 3.* For best effect, brightness and contrast of GFP images were adjusted by setting “minimum” and “maximum” light low and “contrast” high under the “Image” tab.

*Step 4*. If an image was too crowded with worms, the ROI selection tool in the “Analyze” tab was used to crop a sample portion of the stacked images to count. The stacked images were synchronized using Analyze ➔Tools ➔Synchronize windows ➔ Synchronize All commands.

*Step 5*. Worms were counted by eye using the “Multipoint” tool to mark individual worms, with different marking colors for fluorescent and non-fluorescent worms.

### Data Analysis

*i) Mutation Rate*. We calculated the per-generation mutation rate, μ*_i_*, for each MA line *i* as μ*_i_* = *m_i_/n_i_t_i_*, where *m* is the number of mutations, *n* is the number of callable sites, and *t* is the number of generations of MA (Denver et al. 2009). First-order and second-order mutation rates were calculated separately.

Our first question of interest is: does the mutation rate change over time? To test that hypothesis, we can estimate the mutation rates of O1MA lines (µ_0l_) and O2MA lines (µ_02_), calculate the difference Δµ = µ_02_ - µ_0l_, and test the hypothesis that the average difference is significantly different from zero. There is a complication, however, because false inferences affect estimates of µ_0l_ and µ_02_ in different ways (Figure 2).

To proceed, we employed a parametric bootstrap approach. We assume that mutation can be modeled as a Poisson process. We further assume that there is a uniform mutation rate µ = µ_0l_ = µ_02_, i.e., the mutation rate does not change over the course of the experiment.

Because FPs necessarily inflate µ_02_ relative to µ_0l_, the analysis must account for FPs. To begin, we set µ= to the point estimate of µ_0l_. At each generation, an MA line accumulates mutations in its genome drawn from a Poisson distribution with parameter,1_m_ = µL, where µ is the per-generation mutation rate and *L* is the size of the *C. elegans* genome. The total number of mutations in line *i*, m_i_ =,1_m_t_i_, where t_i_ is the number of generations of MA for line *i*. At Gmax=150, the O1MA line is split into the two O2MA sublines, and the mutational process is continued for the remaining Gmax≈150 generations. Each mutation was accepted (“called”) or rejected with binomial probability equal to the proportion of the genome covered in the line (Supplementary Table 2). The number of generations for each MA line was held constant to the observed number for the line. Upon reaching Gmax≈300, *n* FPs were added to each O2MA line, drawn from a Poisson distribution with parameter,1_FP_= the expected number of FPs per line. We next calculate µ_0l_ and µ_02_ from the simulated data and subtract µ_0l_ from µ_02_ to find Δµ. This procedure was repeated 10,000 times to generate a null distribution of Δµ. The value of,1_FP_ for which the observed value of Δµ exceeds 95% of simulated values is the maximum number of FPs per O2MA line that is consistent (at p>0.05) with µ_02_ = µ_0l_, i.e., Δµ > 0 cannot be explained by FPs if µ_02_ = µ_0l_.

The preceding analysis is a hypothesis test. To put confidence intervals on the estimates of µ_0l_ and µ_02_, we used a non-parametric bootstrap, as follows. For µ_0l_, the procedure is:

Step 1. Resample the 96 O1MA lines with replacement, holding the numbers of O1 mutations (*m_i1_*), callable sites (*n_i_*), and O1 generations (*t_i1_*) equal to the observed for each line *i*.

Step 2. To each line, add an additional *FtR* mutations where *FtR* ∼Poisson(λ_FtR_) where λ_FtR_ = 0.022 x *m_i1_*for SNVs and 0.025 x *m_i1_* for indels. These are the mutations which we failed to recall.

Step 3. Calculate the estimated mutation rate µ_0l_ as before, where *m_1_\**=*m_1_*+*FtR*.

Step 4. Repeat the above procedure 10000 times; the upper and lower 2.5% of bootstrap replicates establish the 95% confidence interval on µ_0l_.

For µ_02_, the procedure is:

Step 1. Resample the 192 O2MA lines with replacement, holding the numbers of O2 mutations

*m_i2_*, callable sites (*n_i_*), and O2 generations (*t_i2_*) equal to the observed for each line *i*.

Step 2. For each line *i*, subtract *FP_i_* false positives from *m_i2_*, where *FP_i_* ∼Poisson(λ_FP_) with λ_FP_ = 2 for SNVs and 0.33 for indels. Since the observed number of false positive insertions is zero, but the true probability of a false positive insertion must be non-zero, indels were assigned as deletions or insertions with probability equal to the observed (deletion) bias. .

Step 3. To each line *i*, add an additional *FtR* mutations where *FtR* ∼Poisson(λ_FtR_) where λ_FtR_ = 0.022 x *m_i2_*for SNVs and 0.025 x *m_i2_* for indels. These additional mutations are O2 failutre to recall.

Step 4. Calculate the estimated mutation rate µ_02_ as before, where *m_2_\**=*m_2_-FP+FN*.

Step 5. Repeat the above procedure 10000 times; the upper and lower 2.5% of bootstrap replicates establish the 95% confidence interval on µ_02_.

*ii) Variance in mutation rate*. Also of interest is the question: does the variance in mutation rate (V_µ_) change over time? We address this question in two ways. First, from the same simulations described in the preceding section. For each simulation run we calculate the variance among O1MA lines (V_µ0l_) and O2MA lines (V_µ02_) to generate the relevant null distributions. We then compare the observed variances to the null distributions to test the hypothesis that there is more variance in mutation rate than expected if the mutation rate is uniform (Saxena et al. 2019).

Second, we can – with a caveat - estimate the mutational variance (*V_M_*) for mutation rate. *V_M_*is defined as the per-generation increase in genetic variance for a trait resulting from new mutations and is a fundamental parameter in quantitative genetics (Clayton and Robertson 1955; Lynch and Hill 1986). The among-O1MA family component of variance, *V_L_*, represents the cumulative genetic variance introduced over 150 generations of accumulated mutations . In principle, we would want to estimate *V_M_* for mutation rate from each O2MA line prior to the O2MA phase, because mutations accumulating in the O2 phase will necessarily contribute to the variation among O1MA families after the O2 phase (i.e., at generation 300). That can be easily done in microbes, in which mutation rate can be measured over one round of replication by means of a fluctuation test (Luria and Delbrück 1943), but would require 50% more sequencing in our experiment. Instead, we partition the variance in mutation rate at generation 300 into within- and among-O1MA family components. The among-O1MA family component of variance is *V_L_*; the residual within-family (i.e., between O2MA line) component of variance is *V_E_*.

*V_M_*=*V_L_*/2*t*, where *t* is the number of generations of MA (≈ 150) (Lynch and Walsh 1998, p. 330).

The mutational heritability, 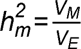. The caveat is that *V_M_* estimated in this way is a lower bound, because of the contribution of O2 mutations to the total variance. Variance components were calculated using restricted maximum likelihood (REML) in a general linear model (GLM) as implemented in the MIXED Procedure of SAS v.9.4, with μ*_O2_* as the dependent variable, O1MA family as a random effect and O2MA line nested within O1MA family as a repeated measure. Statistical significance of *V_M_* was assessed by likelihood ratio test (LRT) of the full model against a model with *V_L_* constrained to zero.

*iii) Fitness effects*. Competitive fitness *W* is defined as the logarithm of the ratio of the frequency of focal strain worms (*p*) to competitor strain worms (1-*p*) on an assay plate, i.e., *W* = log[*p*/(1-*p*)]. To quantify how mutations contribute to competitive fitness, we applied a fully Bayesian mixed-effects modeling framework (Stearns et al. 2025). We compared two Bayesian models: a one-stage (“uniform effects”) model that consolidates all mutations into a single genetic effect structure using a unified genetic covariance matrix, and a two-stage model that explicitly partitions mutations into first-order (O1) and second-order (O2) effects, each with its own genetic parameters and separate covariance matrices. Each value of *W* for observation *i* is modeled as the sum of a global intercept (*c*), a block-specific deviation (*b*_block[*i*]_), genetic effects (*g_i_*), and a residual error (ε*_i_*):

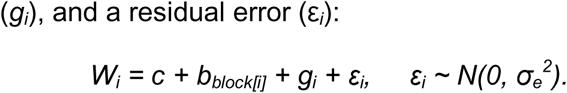

The block effects (*b_j_*) account for systematic batch-to-batch variability arising from separate experimental runs, modeled as independent deviations from a normal distribution:

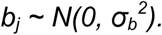

The genetic effects (*g_i_*) capture heritable variation in *W* explained by genetic similarity among strains. Following the Central Limit Theorem, we assume that the cumulative effect of multiple mutations can be approximated by a normal distribution. For each individual *i*, the genetic effect is decomposed into two additive components:

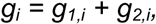

where *g*_1,*i*_ represents the genetic effect from first-order mutations (accumulated from generations 0 to 150) and *g*_2,*i*_ represents the genetic effect from second-order mutations (accumulated from generations 150 to 300).

We assume that the fitness effects of O1 and O2 mutations are drawn from arbitrary DFEs with mean μ*_1_* and μ*_2_*, and standard deviations σ *^2^* and σ *^2^*, respectively. Given that the numbers of observed O1 and O2 mutations are fairly large, assuming mutations combine additively to determine fitness, we can ignore the higher moments of the DFEs and fairly approximate *g_1,i_* and *g_2,i_* using normal distributions by the Central Limit Theorem (see Supplemental Figure 1 in Stearns et al. (2025) for a comparison between the distributions of the sum of mutational effects sampled from arbitrary DFEs and their Gaussian approximations).

Specifically, we sampled the vector of genetic effects (*g_i_*) from a multivariate Gaussian distribution

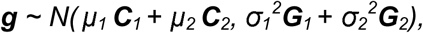

where ***g*** is a vector containing the genetic effects for all observations. ***C****_1_* and ***C****_2_* are diagonal matrices

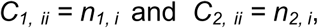

where *n*_1,*i*_ and *n*_2,*i*_ are the number of O1 and O2 mutations in individual *i*. ***G_1_*** and ***G_2_*** account for the covariance in genetic effects between observations, with diagonal terms

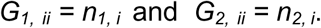

The off-diagonal terms record the number of shared mutations, such that

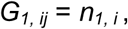

if *i* and *j* share a common ancestor in generation 150, and *G_1, ij_* = 0 otherwise. Similarly,

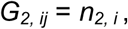

if they share a common ancestor in generation 300.

We fit two competing models under this formulation. In the one-stage (“uniform effects”) model, we assume the O1 and O2 mutations have a common DFE, that is, μ*_1_* = μ*_2_*, and σ*_1_^2^* = σ*_2_^2^*. In the two-stage model, we assume the DFEs of the O1 and O2 mutations are different and independently infer the parameters μ*_1_* and μ*_2_*, σ*_1_^2^* and σ*_2_^2^*.

In addition to one-stage and two-stage model, we also fit a model where we independently estimate the effects of SNVs and indels, while assuming constant DFEs for O1 and O2 mutations. The sampling procedure was a straightforward modification of the steps listed above.

We implemented both models using PyMC (Salvatier et al. 2016) with weakly informative priors to regularize estimation. Intercepts and genetic means were assigned *N*(0,10) priors, while standard deviations received HalfNormal(5) priors. Model parameters were estimated using the No-U-Turn Sampler (NUTS) with 2,000 tuning and 20,000 posterior draws.

To account for potential measurement errors in mutation counts, we implemented a bootstrapping approach that simulates both false positives and false negatives. False positives, which only affect O2 mutations, were modeled using a Poisson distribution with expectation λ = 3. False negatives, which in this context include all mutations missed due to failure to recall, were also modeled using a Poisson distribution with rate

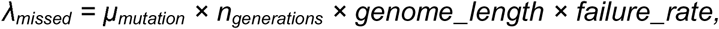

where μ_mutation_ represents the per-generation mutation rate (2.13×10^-9^ for O1, 2.47×10^-9^ for O2), *n*_generations_ = 150, genome_length = 10^8^ bp (the approximate size of the *C. elegans* genome), and failure_rate = 0.025. For each bootstrap iteration, we resampled mutation counts by subtracting false positives from observed O2 counts, adding missed mutations to both O1 and O2 counts, and fitting the Bayesian model with the resampled mutation counts. This approach allows us to propagate uncertainty in mutation counts through to our parameter estimates while maintaining the mathematical structure of our genetic effects model.

## Results

### Mutation rate

Over the first ∼150 generations we observed 2897 putative mutations (2400 SNVs, 341 deletions, and 156 insertions; Supplementary Tables 2 and 7, Supplementary Figure 1). Ignoring false inferences for the moment, the observed first-order per-nucleotide SNV rate µ =1.77x10^-9^/generation; the indel rate µ =0.36x10^-9^/generation. The total first-order mutation rate µ =2.13x10^-9^/generation. In the second ∼150 generations of the experiment we observed 6631 total variants (5031 SNVs, 986 deletions, 614 insertions). The observed second-order SNV rate µ =1.88x10^-9^/generation; the second-order indel rateµ =0.60x10^-9^/generation. The total second-order mutation rate µ = 2.47x10^-9^/generation.

False positives (FP) and false negatives (FN) affect the estimated mutation rates in predictable ways (Figure 2). FPs artificially inflate the inferred µ_02_, whereas FNs do not introduce bias, on average. Because we have only a rough estimate of the number of FPs (≈ 2 SNVs and 0.33 indels per genome; Supplementary Table 3), we tested the null hypothesis that µ_02_ = µ_0l_ by “titrating” the number of FPs by simulation and asking the question: what is the fewest FPs per O2MA line that would lead us to falsely reject the null hypothesis µ_02_ = µ_0l_ (i.e., Type I error)? For SNVs, there must be fewer than 0.6 FPs for the observed difference Δµ_SNV_ = 0.23x10^-9^/generation to not be sufficiently explained by FPs (Figure 3A), so we conclude that there is no evidence that µ_02,SNV_ > µ_0l,SNV_ . For indels, the same test shows the critical number of FPs per line is 2.71 (Figure 3B). The probability of observing no more than one FP in three replicates (see Methods and Supplementary Table 3) if the true Poisson mean is 2.71 is approximately 0.0028, so we conclude that the evidence suggests that µ_02,Indel_ > µ_0l,Indel_.

**Figure 3.**
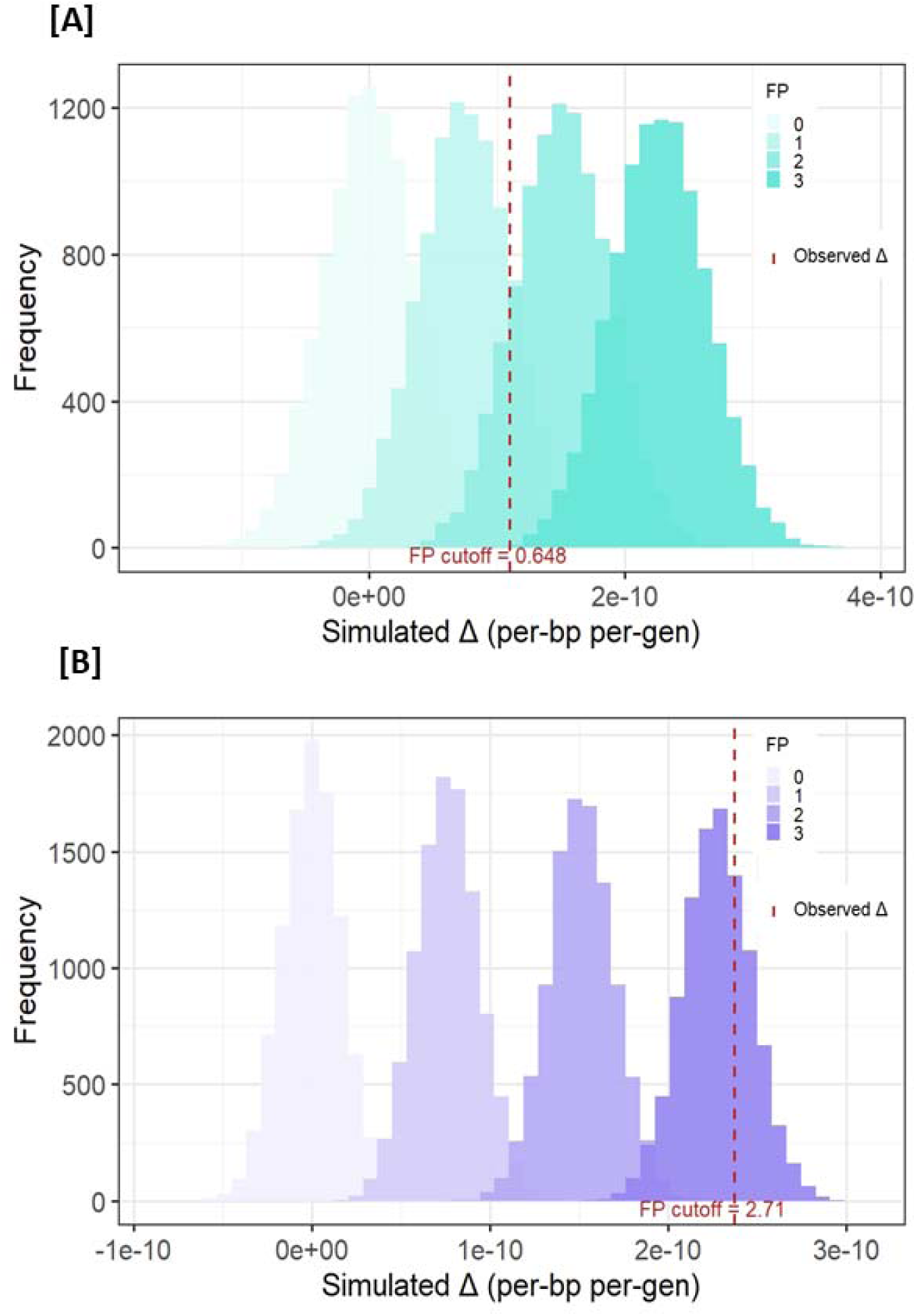
Distribution of simulated values of Δμ*=*μ*_O2_-*μ*_O1_* with increasing number of false positives. Red dashed line is the observed value. “FP cutoff” is the minimum number of false positives necessary for the data to be consistent with the hypothesis Δμ=0. (**A**) SNVs (**B**) Indels.

Estimates of mutation rates corrected for false inferences and 95% confidence intervals are given in Table 1.

**Table 1.** Summary statistics. *N_Lines_* is the number of MA lines, is the average number of generations of MA (top row) and number of days of MA (bottom row). Values of µ are mutation rates (><10^9^) calculated from the observed number of mutations without correction for false inferences. Values of µ^ are estimates of mutation rates (><10^9^) including correction for false inferences at the 3X coverage criterion. Estimated mutation rates (µ^) are calculated as 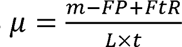, where *m* is the observed number of variants, *FP* is the number of false positives, *FtR* is the number of mutations failed to recall and *L* is the number of bases called at 3X coverage. See Methods for details of the calculations.

| | $N_{Lines}$ | %Genome<br>(SEM) | $\bar{t}$<br>(SEM) | $\bar{\mu}_{SNV}$<br>(SEM) | $\bar{\mu}_{DEL}$<br>(SEM) | $\bar{\mu}_{INS}$<br>(SEM) | $\bar{\mu}_{TOTAL}$<br>(SEM) | $\hat{\mu}_{SNV}$<br>(95% CI) | $\hat{\mu}_{DEL}$<br>(95% CI) | $\hat{\mu}_{INS}$<br>(95% CI) | $\hat{\mu}_{TOTAL}$<br>(95% CI) |
| --- | --- | --- | --- | --- | --- | --- | --- | --- | --- | --- | --- |
| O1 | 96 | 97.25<br>(0.051) | 145.9<br>(0.8) | 1.77<br>(0.027) | 0.25<br>(0.011) | 0.113<br>(0.0071) | 2.13<br>(0.030) | 1.78<br>(1.73-1.84) | 0.25<br>(0.23-0.27) | 0.12<br>(0.10-0.13) | 2.15<br>(2.09-2.21) |
|  |  |  | 600 | 0.427<br>(0.0062) | 0.061<br>(0.0027) | 0.028<br>(0.0017) | 0.52<br>(0.0072) | 0.43<br>(0.42-0.44) | 0.062<br>(0.056-0.067) | 0.028<br>(0.025-0.032) | 0.52<br>(0.508-0.536) |
| O2 | 192 | 97.25<br>(0.051) | 144.8<br>(1.6) | 1.88<br>(0.032) | 0.37<br>(0.016) | 0.23<br>(0.013) | 2.47<br>(0.046) | 1.75<br>(1.68-1.81) | 0.36<br>(0.33-0.39) | 0.22<br>(0.20-0.25) | 2.33<br>(2.24-2.42) |
|  |  |  | 624 | 0.43<br>(0.0075) | 0.084<br>(0.0034) | 0.053<br>(0.0026) | 0.57<br>(0.0098) | 0.40<br>(0.39-0.42) | 0.082<br>(0.075-0.087) | 0.051<br>(0.046-0.056) | 0.54<br>(0.52-0.56) |

Some types of mutations occur independently of genome replication and accrue at an approximately constant rate in chronological time (Gao et al. 2016; Gao et al. 2019). If the chronological mutation rate remains constant but the generation time increases, the result will be an increase in the per-generation mutation rate. There was little difference in the average number of generations between the O1 and O2 lines over the course of the experiment (O1 mean = 145.9 generations; O2 mean = 144.8 generations), and the conclusions drawn from mutation rates scaled per-generation apply similarly to mutation rates scaled per-day (Table 1). The number of indels was positively correlated with generations of MA in both O1 (*r_O1,indel_*=0.29, p<0.004) and O2 lines (*r_O2,indel_*=0.25, p<0.001). In the O1 lines, the number of SNVs was uncorrelated with the number of generations (*r_O1,SNV_*=0.10, p>0.32), whereas in the O2 lines the number of SNVs was positively correlated with the number of generations (*r_O2,SNV_*=0.46,p<<0.0001). The lack of correlation of number of SNVs in the O1MA lines is likely due to lack of power resulting from the smaller sample size and the lack of variation in generations of MA. Taken together, we reject the hypothesis that mutation rate is independent of generation time, but we cannot rule out a contribution of replication-independent mutation.

### Mutation spectrum

Consistent with the observed lack of evolution of µ_SNV_, the SNV mutation spectra did not differ between O1MA and O2MA lines (Monte Carlo Fisher’s Exact Test, p > 0.12) (Figure 4). G:C→A:T transitions (28.6%) and A:T→T:A transversions (28.1%) were the most common classes of base substitutions, as is typical in *C. elegans* MA experiments (Rajaei et al. 2021; Konrad et al. 2019).

**Figure 4.**
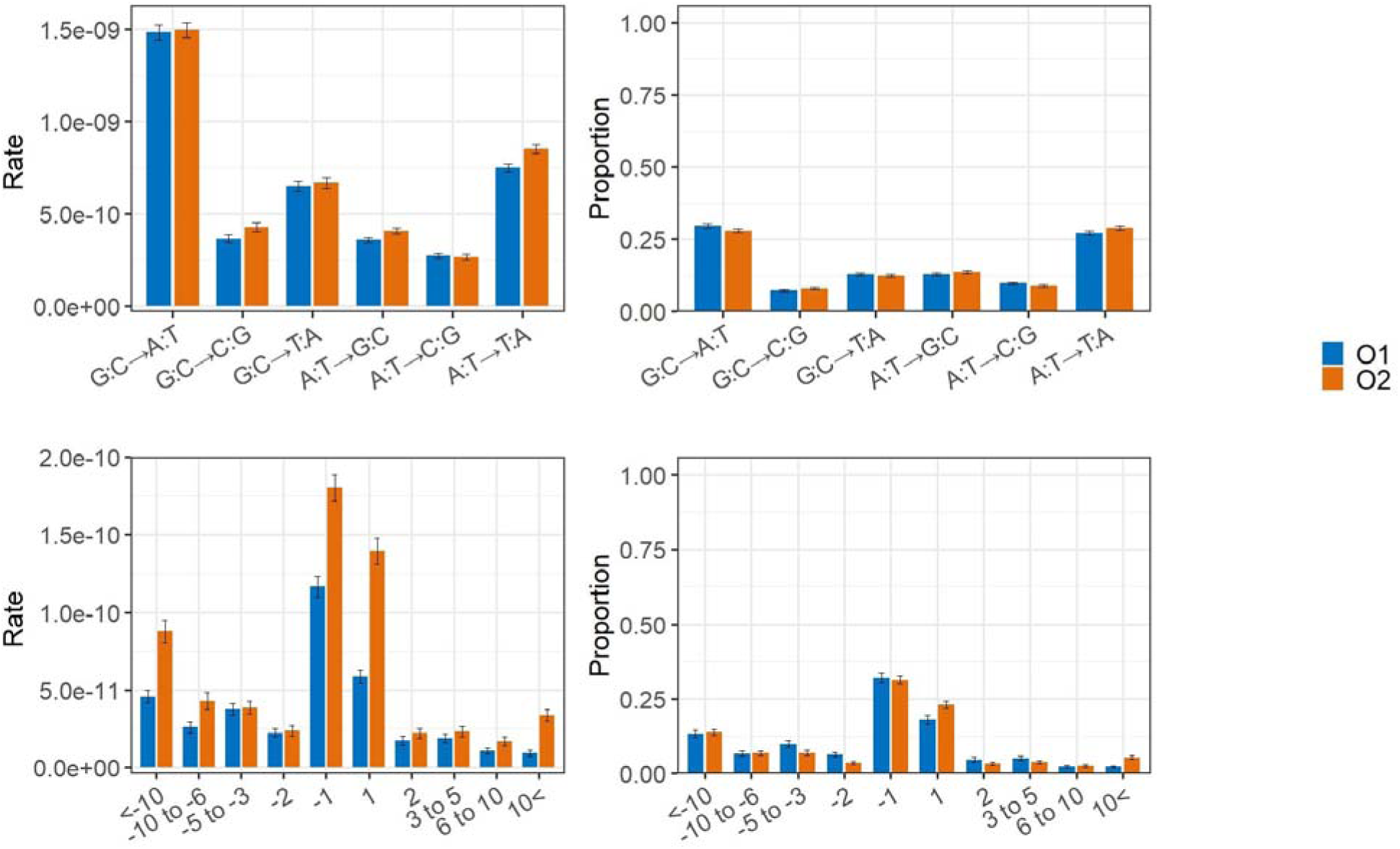
Type-specific mutation rates and spectra. (**Top left**) SNV mutation rates (**Top right**) SNV frequency spectrum **(Bottom left**) Indel mutation rates (**Bottom right**) Indel frequency spectrum. Error bars are SEM of MA lines.

The indel spectrum, in contrast, was significantly different between O1MA and O2MA lines (Figure 4). In the O1MA lines, deletions outnumbered insertions by nearly 2.2:1, whereas the deletion bias in the O2MA lines was significantly reduced to ∼1.6:1 (Fisher’s Exact Test (FET), p < 0.0001). Considered separately, the deletion spectrum differed significantly between O1 and O2 lines (FET, p < 0.003), with an increase in the shortest (1 bp) and longest (>10 bp) categories and concomitant decreases in the intermediate categories. The insertion spectrum also differed between O1 and O2 (FET, p < 0.0001), also with increases in the 1 bp and >10 bp categories and decreases in the intermediate categories.

### Multi-nucleotide variants (MNVs)

It is known that mutations occur in clusters more often than expected by chance (Schrider et al. 2011), and that short tandem repeats tend to accumulate SNVs (“imperfect microsatellites”) in addition to indels (Saxena et al. 2019). We (arbitrarily) define an MNV cluster as all variants within 100 bp of their nearest neighbor, although >90% of all MNV mutations occur within 20 bp of their nearest neighbor. To quantify the rate and spectrum of MNV mutations and to account for the possibility that MNVs may result in mapping errors that incorrectly identify one O1 MNV as two different O2 mutations, one in each O2 MA sibling line, we re-analyzed the data with and without MNVs (Supplementary Table 8). At the liberal 3X coverage criterion, more O2 SNVs occur in MNVs than O1 SNVs (7.2% O2 vs. 4.4% O1; Pearson’s chi-square = 20.3, P<0.0001). In contrast, the proportion of insertions in MNVs is greater in O1 than O2 (34.0% O1, 22.5% O2; Pearson’s chi-square = 8.82, P<0.01). The proportion of deletions in MNVs does not differ significantly between O1 and O2 (16.2% O2 vs 12.6% O1; Pearson’s chi-square = 2.56, P>0.10). The results are similar at the more stringent 10X coverage criterion (Supplementary Table 8). Summary statistics of mutation rate with MNVs considered as a separate class of mutation are given in Supplementary Table 9.

To account for the possibility that O1 mutations in MNVs may be mis-mapped as two different O2 mutations, we re-ran the FP “titration” analysis described above with mutations in MNVs omitted. The conclusions do not change: the SNV rate does not change significantly from O1 to O2, whereas the indel rate increases by about 60% from O1 to O2 (critical FP = 2.24, Poisson P=0.009; Supplementary Figure 2). The greater proportion of SNVs in MNVs from O1 to O2 can be explained by the increased rate of indel mutations (especially insertions) from O1 to O2 if the mutational process that generates indels also tends to generate associated SNVs.

### Variance in mutation rate

To begin, we ask: can the observed variance in mutation rate among O1MA lines (V_µ0l_) be sufficiently explained by sampling variance of a uniform mutational process? For SNVs the answer is yes; V_µ0l,SNV_ is sufficiently explained by sampling if mutation conforms to a uniform Poisson process (p>0.09, Supplementary Figure 3A). For indels, however, V_µ0l,Indel_ is too great to be explained by a uniform Poisson process (p<0.02, Supplementary Figure 3B).

To extend the analysis to the O2MA lines, FPs must be accounted for. We assume an average of 2 SNV and 0.33 indel FPs per sample. For both SNVs and indels there is more variance among O2MA lines than is consistent with a uniform mutational process (Supplementary Figure 3).

Estimates of mutational variances (V_M_) for mutation rate do not differ significantly from zero for SNVs, indels, or total mutation rate (Supplementary Table 10). REML estimates of the among-O1MA family component of variance (V_L_) for SNVs and total mutation rate are 0; the point-estimate of the mutational heritability (h_m_) for indel rate is on the order of 10 /generation, roughly an order of magnitude less than for typical quantitative traits (Houle et al. 1996; Conradsen et al. 2022). The lack of significant V_M_ for SNV mutation rate is consistent with the inferred uniformity of the O1MA process. For indels, the simplest explanation for the discrepancy between the two ways of quantifying variance (V_µ0l_ vs. V_M_) is that the effects of mutations accumulated in the O2 phase largely swamp those of the O1 mutations, removing the signal of heritability (although when the strong mutator O1MA family 426 is included in the analysis, h_m_ for indel rate ≈ 0.1/generation, which we take as a cautionary note).

### Mutational effects on fitness

Because all mutations in an MA line are in complete linkage disequilibrium, the effects of individual mutations are unidentifiable. However, knowing the number of mutations in each MA line allows a straightforward estimate of the first two moments of the DFE, because the sum of the effects of a large number of individual mutations within a line can be well-approximated by a normal distribution regardless of the underlying DFE, according to the Central Limit Theorem (see Methods for explanation).

Competitive fitness (*W*) decreased by about 0.2% per generation over the first 150 generations of MA and by about half that over the next 150 generations of MA (Supplementary Figure 4, Supplementary Tables 11 and 12). There is a weak negative association between mutation rate and fitness (Supplementary Figure 5).

Given that the number of mutations is similar between O1MA and O2MA lines, the implication is that the mean effect on competitive fitness of O1 mutations (*u_O1_*) is greater than that of O2 mutations (*u_O2_*). To test that hypothesis, we compared a model in which the mean mutational effect *u* is constrained to remain constant over the entire 300 generations (“uniform effects”) to a “two-stage” model in which *u_O1_* and *u_O2_* are allowed to differ. Models were compared using the Widely Applicable Information Criterion (WAIC) and leave-one-out cross-validation (LOO) to assess model fit and predictive performance (Vehtari et al. 2017). The differences between models are substantial (Δ_WAIC_= 13.52; Δ_LOO_ = 12.69) and favor the uniform effects model. The point estimate of the mean mutational effect *u* = -0.0139, with variance σ*_g_*^2^ = 1.22x10^-4^ (Figure 5).

**Figure 5.**
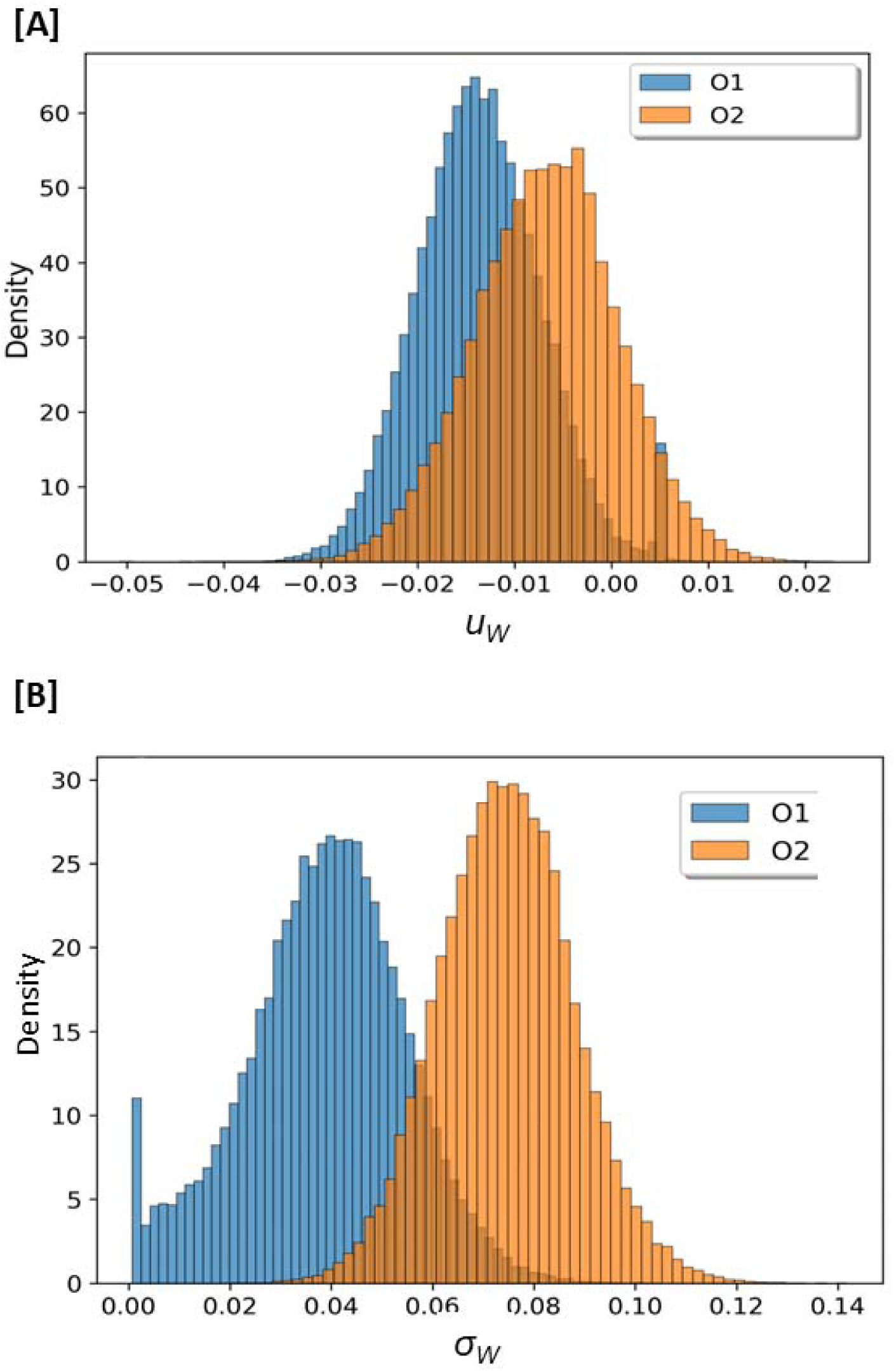
Distribution of posterior probabilities of mutational effects *u* on competitive fitness. (**A**) Mean *u* (**B**) Standard deviation σ*_G_*.

Given the different mutational properties of SNVs and indels, we attempted to assess potential differences in fitness effects between the two types of mutations by fitting a model allowing separate DFE parameters for the two mutation classes. However, the WAIC values for the one-DFE and two-DFE models are virtually identical, providing no evidence for different fitness effects between SNVs and indels. Although it is possible that the average fitness effects of SNVs and indels do not differ, the result is more simply explained by the limited statistical power resulting from the smaller number of mutations in each class.

The O2MA design permits us to address the question: is the O2 mutation rate a function of O1 fitness? The absence of significant mutational heritability for mutation rate does not preclude *a priori* a relationship between O1 fitness and O2 mutation rate, unless O1 mutation rate strongly predicts O1 fitness, which it does not (Supplementary figure 5). However, O2 mutation rate (SNV, indel, and total mutation rate) is not correlated with O1 fitness (Pearson’s *r*<0.09, p>0.59 in all cases.

## Discussion

Our foremost interest is in the evolution of the mutation rate. Mutator alleles (e.g., a defective allele of a DNA repair gene) increase the mutation rate; antimutator alleles (e.g., an allele at a regulatory sequence that increases the expression level of a DNA repair gene) decrease the mutation rate. Intuition suggests that spontaneous mutator mutations should be more common than antimutator mutations, on the grounds that biological systems are well-adapted and a random change is more likely to make a well-adapted system worse than it is to improve it (Drake 1993; Fisher 1930; Lynch 2012). That intuition leads to a clear prediction: if spontaneous mutations are allowed to accumulate unimpeded by natural selection, the mutation rate should increase over time. However, one can imagine alternative scenarios in which the converse is true. For example, if (*i*) mutations accrue approximately constantly in chronological time and (*ii*) growth rate (and thus generation time) is a function of metabolic rate and (*iii*) deleterious mutations decrease metabolic rate on average, then a random mutation might be expected to decrease the per-generation mutation rate. Or along the same lines, if metabolic byproducts (e.g., free radicals) are an important cause of mutation (Martin and Palumbi 1993; Gillooly et al. 2005) and if deleterious mutations decrease metabolic rate on average, the mutation rate might decrease as deleterious mutations accumulate.

Given the importance of mutation to all areas of biology, and the vast trove of theoretical investigations of the evolution of mutation rate (Drake et al. 1998; Lynch et al. 2016), there have been surprisingly few empirical studies that have attempted to quantify the effects of spontaneous mutations themselves on the rate and spectrum of mutation. In the first such study of which we are aware, Ávila et al. (2006) initiated an O2MA experiment from a single high-fitness MA line of *Drosophila melanogaster*. Their study predated economical whole-genome sequencing, but they found that the decline in fitness in the O2MA experiment was greater than in the O1MA experiment. They attributed the increased rate of decline in fitness to an increase in mutation rate, although they could not formally rule out greater mutational effects in the O2MA experiment (i.e., synergistic epistasis).

Sharp and Agrawal (2016) compared the spontaneous mutation rates between a wild-type strain of *D. melanogaster* and derivative strains engineered to carry mutations of large effect, to test the hypothesis that genomes that carry a high load of deleterious mutations mutate faster than less-loaded genomes (Agrawal 2002; Shaw and Baer 2011).. They found that mutation rate of the mutant strains was higher, due to an increased frequency of short deletions, which they attributed to differences in the repair of double-strand breaks.

Saxena et al. (2019) did an O2MA experiment with *C. elegans*, also designed to investigate the relationship between initial mutation load and mutation rate. The indel rate (specifically the insertion rate) increased significantly in the O2MA lines; contrary to expectation, however, the O2MA indel rate was greater in the high-fitness (= less-loaded) lines. The SNV rate increased on average, but not significantly. Two of the ten O1MA families (n=5 O2MA lines per O1MA family) evolved SNV O2 mutation rates that were significantly different from the ancestor; one family increased and the other decreased.

Liu and Zhang (2021) brought the power of yeast genetics to bear on the question of mutation rate evolution. They first constructed a set of O1MA lines in which a mismatch repair gene (*MSH-2*) was deleted by CRISPR-Cas9 engineering to increase the mutation rate. They then constructed a set of O2MA lines by re-inserting the *MSH-2* gene into the O1MA lines and allowing mutations to accumulate further. They found that the O2MA mutation rate increased on average, but a surprisingly high proportion of the O2MA lines (12/48) evolved a decreased mutation rate. Moreover, they identified a gene with antagonistic pleiotropic effects on cellular physiology (autophagy) and mutation rate.

Classical theory predicts that, in the presence of recombination, direct selection favors a reduction in the mutation rate except under special circumstances (an example of the “reduction principle”, Liberman and Feldman 1986), which leads directly to “Sturtevant’s Paradox”: why does the mutation rate not evolve to zero? (Sturtevant 1937). There are two competing (but not mutually exclusive) hypotheses for why the mutation rate does not evolve to zero. In historical chronological order, the Cost of Fidelity (CoF) hypothesis (Supplementary Figure 6A) posits that the mutation rate is under stabilizing selection, with the optimum established by a tradeoff between direct selection to reduce the input of deleterious mutations and indirect selection to minimize the resources (time, energy, molecules) devoted to replication fidelity and genome surveillance (Kimura 1967). At an equilibrium mutation rate established by the CoF, the per-generation fitness cost of an antimutator must equal the cumulative per-generation fitness cost of the deleterious mutations spawned by an equivalent mutator.

In contrast, the Drift Barrier hypothesis (Supplementary Figure 6B) posits that the mutation rate is always under directional selection downwards to reduce the input of deleterious mutations (per the reduction principle), but that at some point the fitness increase resulting from a further reduction of the mutation rate is less than the reciprocal of the effective population size (*N_e_*), at which point selection is too weak to counteract genetic drift (Lynch 2008, 2011).

The existence of a mutational bias means that the trait is under directional selection in the direction opposed to the bias in the vicinity of the ancestral (i.e., unmutated) mean (Charlesworth 2013a; Lynch and Menor 2025). However, there is no information about the fitness function outside the range of the data. A mutational bias, as we observed for indels, fulfills a strong prediction of the DB hypothesis, but does not refute the existence of a non-zero fitness optimum, as predicted by the CoF. In contrast, an increase in the trait variance in the absence of a mutational bias, as we observed for SNVs, is not consistent with the DB hypothesis, and fulfills a key prediction of the CoF.

Our findings about the evolution of the mutation spectrum reinforce and extend previous findings from *C. elegans* and other organisms, in which various types of physiological stress – exogenous and endogenous – increase the indel mutation rate, but have little if any effect on the base substitution mutation rate. Rajaei et al. (2021) performed an MA experiment with an N2-derived strain of *C. elegans* carrying a mutation (*Mev-1*) that causes increased cellular oxidative stress. In that experiment, the indel (and specifically, the insertion) rate increased by about 50% over wild-type, whereas the SNV mutation rate decreased slightly. As noted above, MA lines of *D. melanogaster* initiated from strains engineered to carry large-effect deleterious mutations experience a higher rate of short deletions than their wild-type controls, whereas the SNV mutation rates do not differ (Sharp and Agrawal 2016). *Arabidopsis thaliana* MA lines propagated under benign and stressful conditions of salinity (Jiang et al. 2014) and temperature (Belfield et al. 2020) show large and highly significant increases in indel mutation rates compared to much smaller increases in SNV rates. A similar increase in indel frequency was reported for the green alga *Chlamydomonas reinhardtii* MA lines maintained under salinity stress (Hasan et al. 2022). Several studies have investigated the mutational properties of *C. elegans* in MA experiments done with elements of the DNA repair system knocked out, which has the overall effect of greatly increasing the indel rate, but the bias is typically toward deletions rather than insertions (Meier et al. 2014; Meier et al. 2018; Volkova et al. 2020).

Our second main question of interest concerns the evolution of mutational effects on fitness. Thirty years ago, Kondrashov (1995) famously posed the question “Why have we not died 100 times over?”. Given the size of the human genome (large) and effective population size (much smaller), the continued existence of humans seemed incompatible with the inferred genetic load (∼100 lethal equivalents) if the effects of deleterious mutations are independent. The simplest explanation is that the effects of deleterious mutations are not independent, but rather they interact synergistically, i.e., epistasis is negative. However, it is not immediately obvious why deleterious mutations should interact synergistically on average. One possible explanation is that selection is “soft” (Wallace 1968), wherein population absolute fitness is established by resource availability and relative fitness results from competition among individuals for resources (Peck and Waxman 2000; Agrawal and Whitlock 2012). If so, it implies that epistasis is more likely to be detected under competitive conditions, as we report here.

This study provides no support for the possibility that deleterious mutations interact synergistically, and in fact the point estimate of the effect of second-order mutations (*u_O2_*) is about half that of the first-order effect, suggestive of diminishing returns epistasis. However, we can confidently reject the two-stage model of mutational effects in favor of the one-stage model of constant mutational effects (Δ_WAIC_ ≈ 12). There are obviously many caveats associated with that inference, including environmental context and statistical power. We note that our findings recapitulate those of Peters and Keightley (2000) and Katju et al. (2015) who looked for epistatic effects of new mutations on non-competitive fitness in *C. elegans* and did not find them.

*Conclusions and Future Directions*. The results of this study are clear: there is an upward mutational bias for indels over the course of 300 generations, whereas there is no such bias for SNV mutations, although the variance in µ_SNV_ increased over the course of the experiment. The discrepancy between evolution of the indel and SNV mutational processes, in this study and others, is intriguing. One possibility is that the genomic mutational target for the control of the indel process is simply larger than that of SNVs, and that if selection was relaxed for long enough that the SNV mutation rate would begin to drift upward. We suspect this is probably true. However, it is worth considering the possibility that the indel and SNV mutational processes are subject to different patterns of selection, with the indel rate under directional selection downward (the DB) and the SNV rate under stabilizing selection (the CoF). We are unaware of any study in which any element of the mutation rate has drifted significantly downward.

What remains to be determined is the nature of the fitness function on the downward (antimutator) side of the ancestral mean mutation rate. The obvious way to address that question is to assess fitness as a function of mutation rate over the course of one generation, from a source population with significant variation in mutation rate. That can be done straightforwardly in microbes by means of a fluctuation test. Liu and Zhang (2021) did exactly that and reported that mutation rate of their yeast O2MA lines is uncorrelated with growth rate. Unfortunately, multicellular organisms do not lend themselves to fluctuation tests, and a suitably powered one-generation experiment using our methods would be extremely labor intensive and require a great deal of sequencing. However, genome sequencing is getting ever cheaper, and advances in high-throughput phenotyping are promising (O’Brien et al. 2025; Mok et al. 2023; Tintori et al. 2025).

The idea that epistasis among deleterious mutations should be synergistic on average has been as influential among evolutionary geneticists as the supporting evidence for it is weak (de Visser et al. 2011). One obvious possibility is that 50+ years of inconclusive evidence is the result of underpowered experiments intended to quantify an inherently noisy trait. It is now possible to disentangle the confounding between mutation rate and mutational effects that bedeviled early studies (e.g., Ávila et al. 2006; Mukai 1969), and here too, high-throughput phenotyping holds promise. However, the intriguing possibility exists that epistasis itself is not only evolvable (Azevedo et al. 2006; Desai et al. 2007), but that the direction of epistasis favored by selection is dependent on idiosyncratic properties of the population including effective population size and mutation rate (Sydykova et al. 2020; Gros et al. 2009). The truth may already be out there.

## Data Availability

Raw sequence data are deposited in the NCBI Sequence Read Archive BioProject #PRJNA1089216. All data in the Supplementary Tables are also deposited in Figshare at https://doi.org/10.6084/m9.figshare.30197935. Code and unprocessed image files are available at https://github.com/Baer-Group/O2MAno2.

## Acknowledgments

David Julian generously provided access to his microscope for the fitness assays, and Tim Crombie assisted with microscope software. Ezgi Karabulut, Eileen Kelly, and Hope Kerr assisted with counting worms. We thank the reviewers and the Associate Editor for their many insightful comments and suggestions. Support was provided by National Institutes of Health awards GM127433 (CFB, VK, and ECA) and GM154908 (JZ).

