## Supplementary Figures 1-6 for "Evolution of the rate, molecular spectrum, and fitness effects of mutation under minimal selection in *Caenorhabditis elegans*"

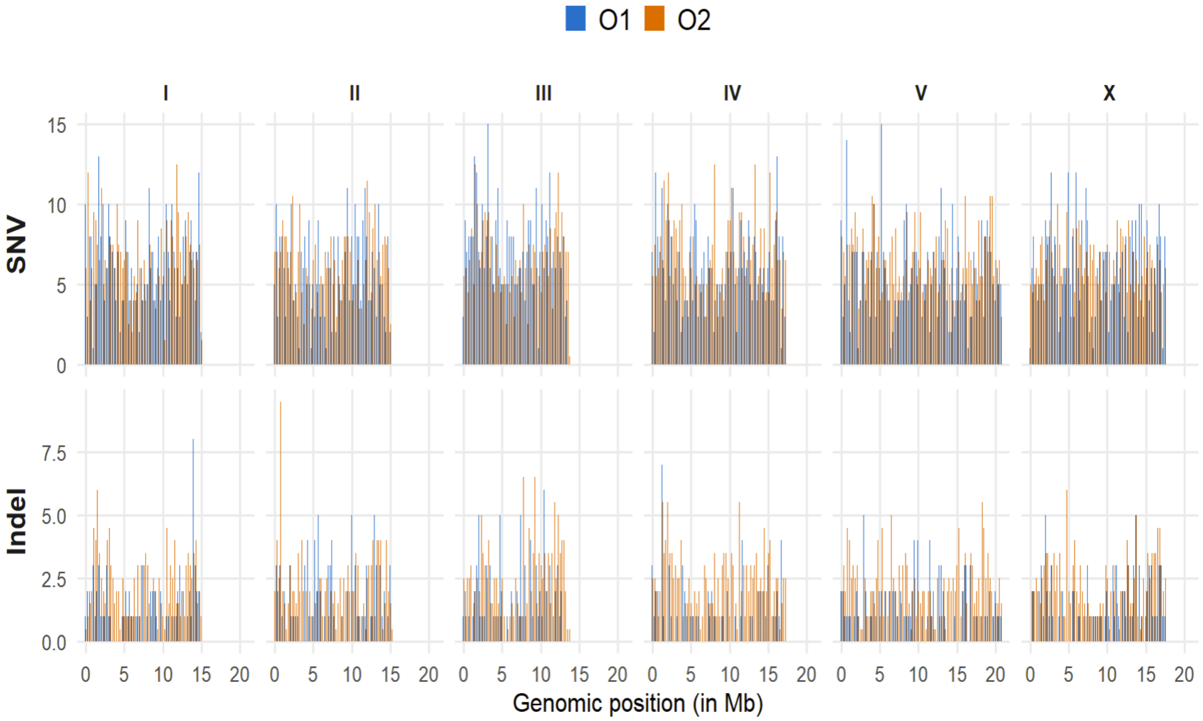


**Supplementary Figure 1**. Distribution of mutations along chromosomes. Top panel, SNVs; bottom panel, indels.

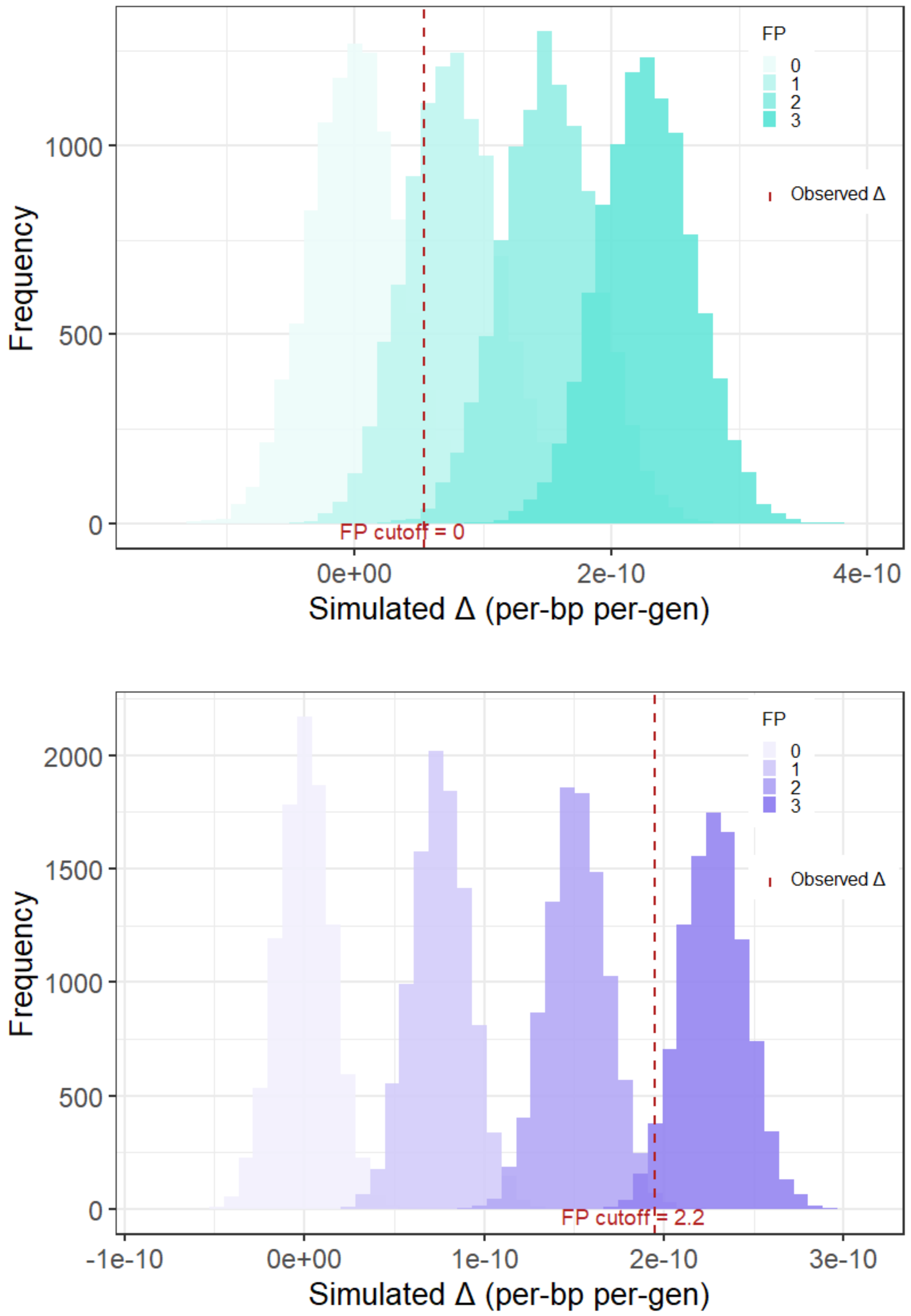


**Supplementary Figure 2**. Distribution of simulated values of $\Delta\mu=\mu_{O2}-\mu_{O1}$ with increasing number of false positives. Variants in MNVs are omitted from the analysis. Red dashed line is the observed value. “FP cutoff” is the minimum number of false positives necessary for the data to be consistent with the hypothesis $\Delta\mu=0$. (**A**) SNVs (**B**) Indels.

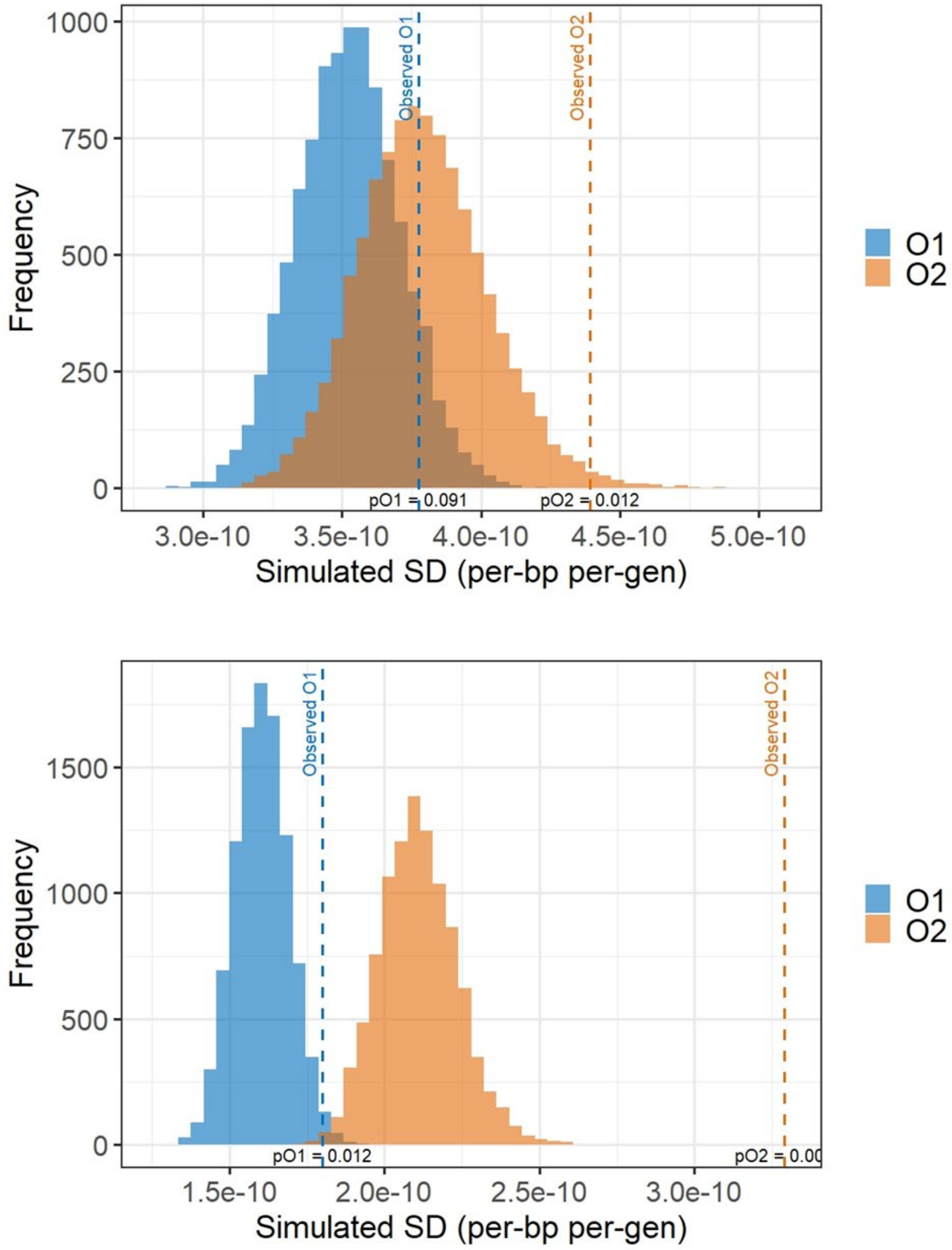


**Supplementary Figure 3**. Simulated distributions of variance in line mean mutation rates under the assumption of uniform mutation rate and the observed average number of false positives (FP) per line. Vertical dashed lines show the observed values. (**Top**) SNVs, 2 FPs/line (**Bottom**) Indels, 0.33 FPs/line.

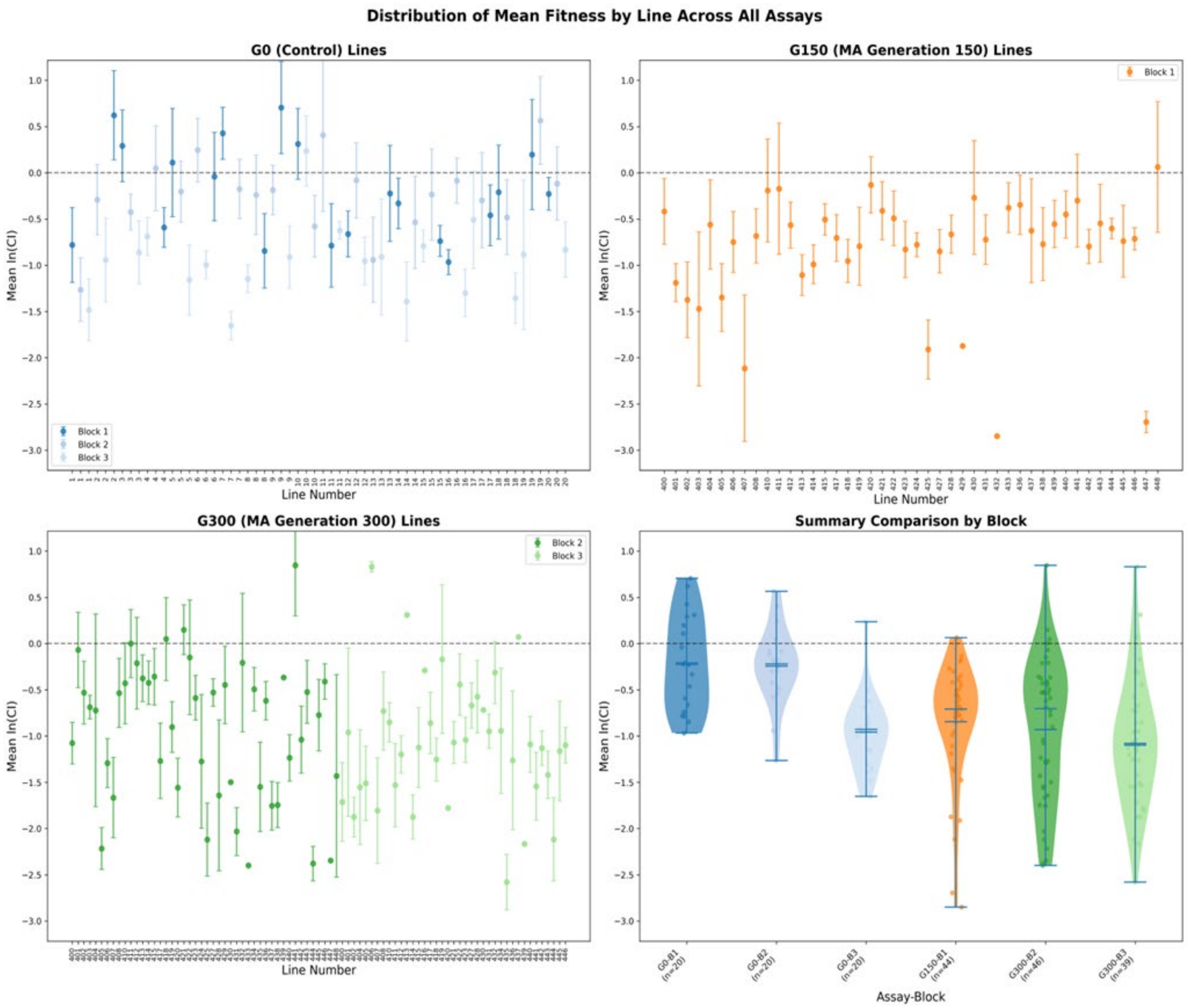


**Supplementary Figure 4**. Distribution of line mean competitive fitness *W* by treatment (MA vs G0) and assay block. (**A, top left**) G0 pseudolines (**B, top right**) G150 MA lines (**C, bottom left**) G300 MA lines (**D, bottom right**) Comparison across assay blocks.

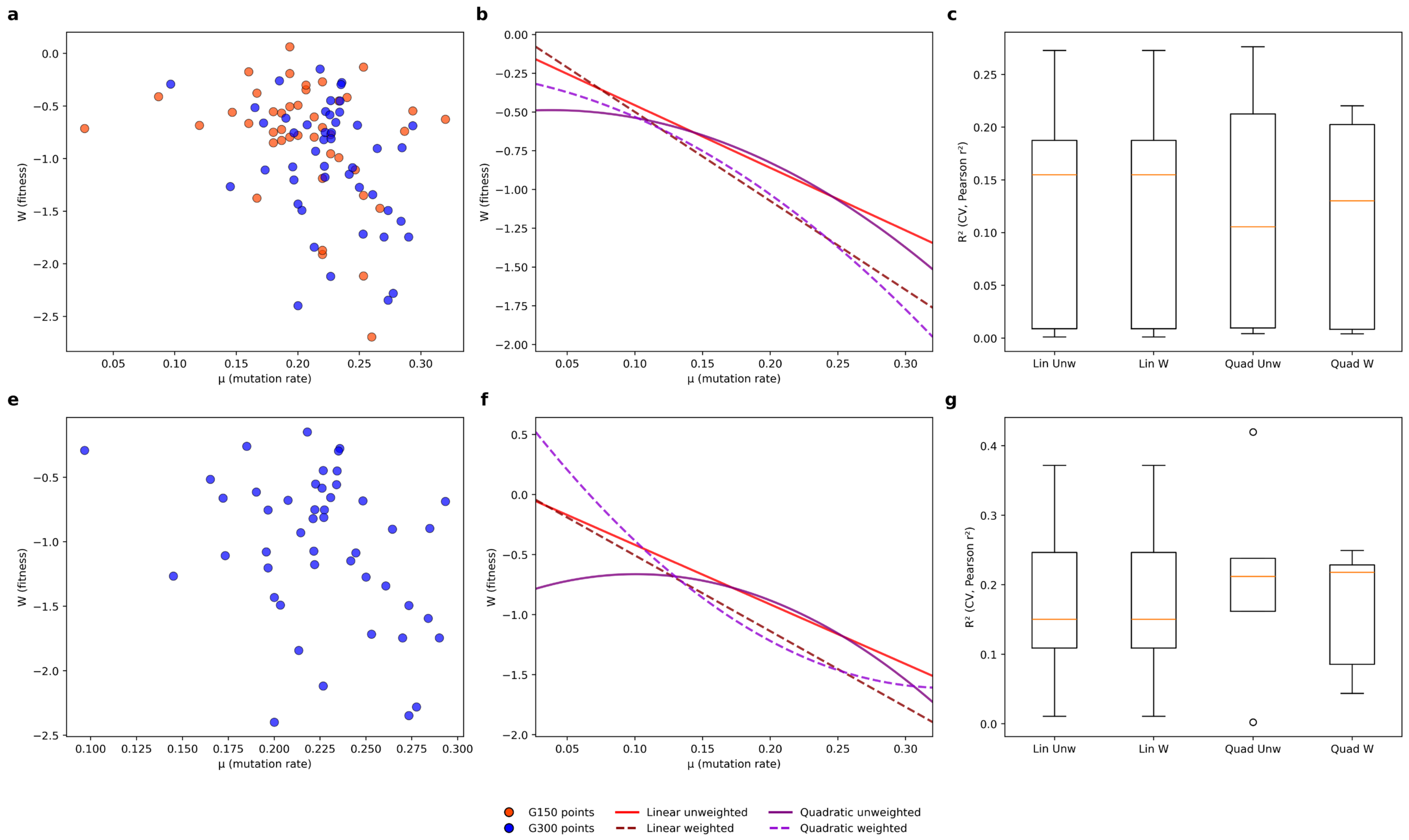


**Supplementary Figure 5**. Regression of competitive fitness (*W*) on total mutation rate (*μ*). (**a-c**) All data (**e-g**) Generation 300 O2MA lines only. Orange lines in panels **c** and **g** are medians.

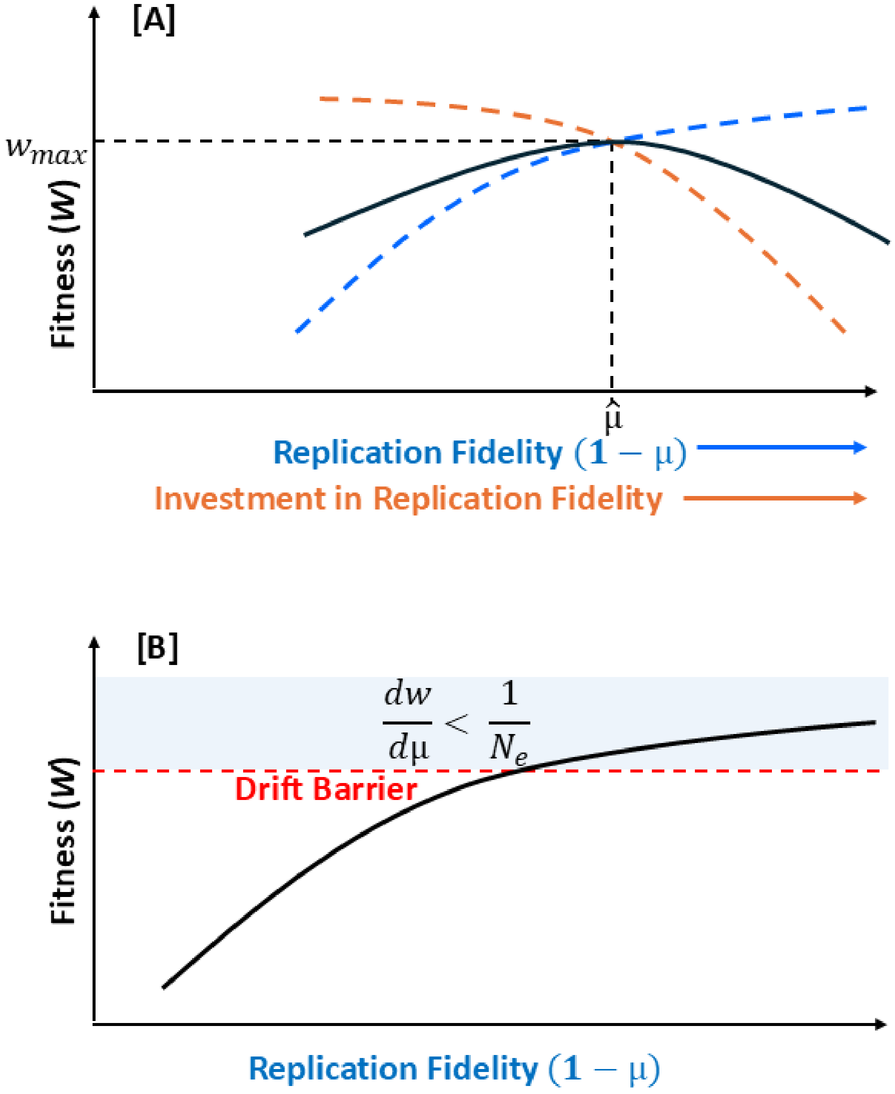


**Supplementary Figure 6**. Theoretical models of mutation rate evolution. $\mu$ represents mutation rate; replication is perfect when $\mu$ = 0. (**A**) Cost of Fidelity. (**B**) Drift Barrier.
