## Supplementary Appendix 1 for "Evolution of the rate, molecular spectrum, and fitness effects of mutation under minimal selection in *Caenorhabditis elegans*"

### *How to assay MA-line worms with Image-J*

Step 1: Start ImageJ. Click and drag image files to taskbar to open

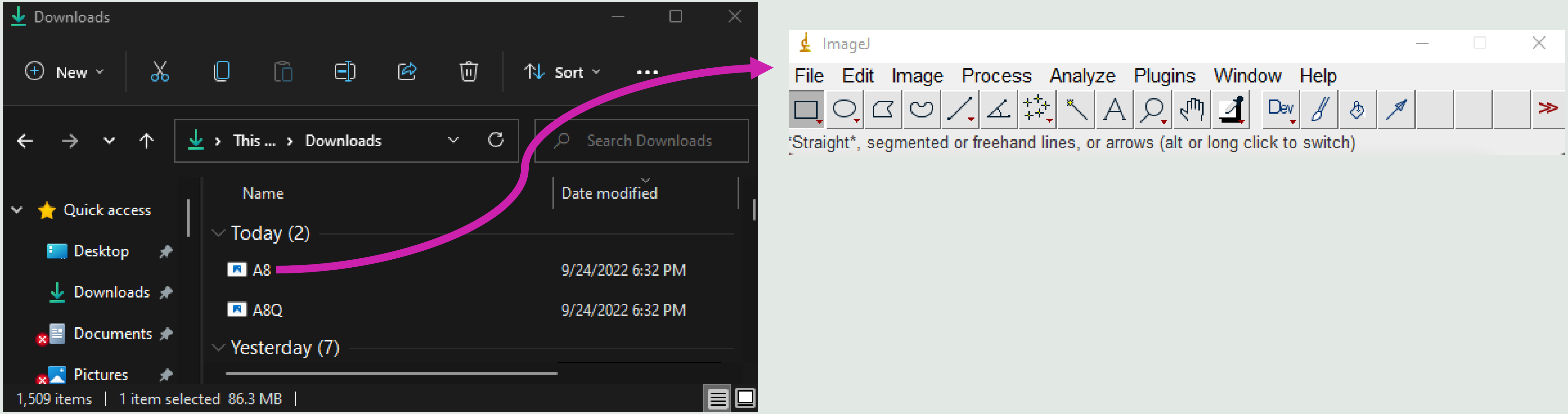

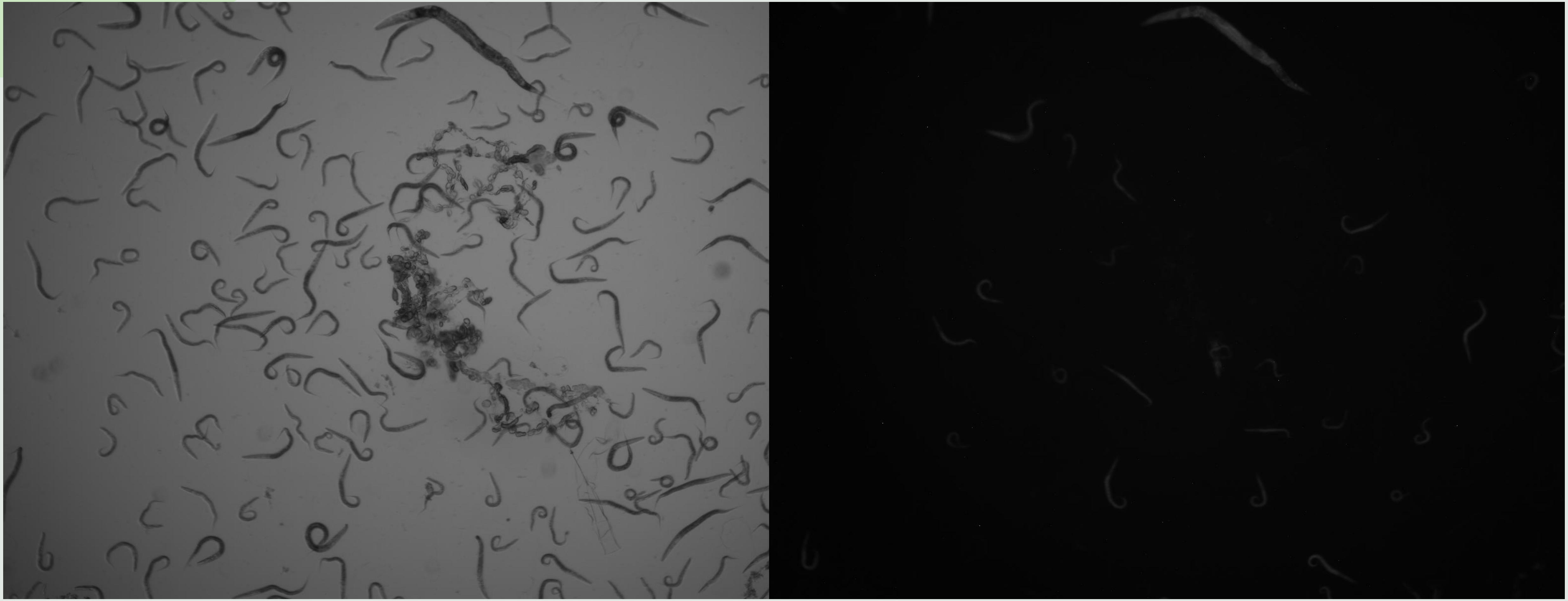

Bright-field image: Both MA-line and N2 Bristol worms are visible

GFP fluorescent image: Only MA-line mutants visible

### Step 2: Adjust brightness and contrast settings of GFP images

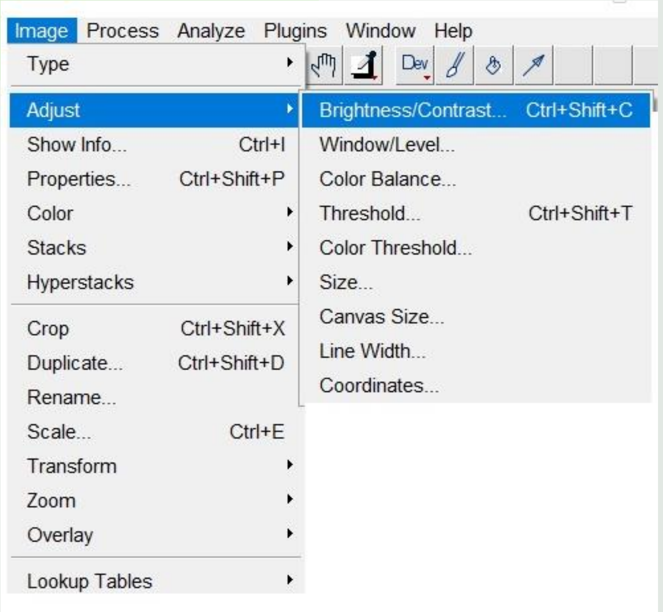


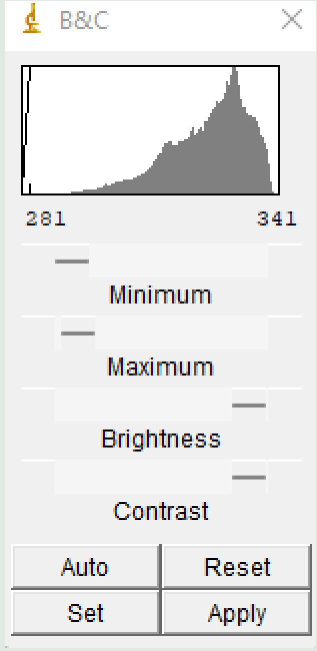


For best effect, set minimum and maximum light settings low and contrast high. Adjust to make sure all GFP-labeled worms in are still visible. Click “Apply” when done.

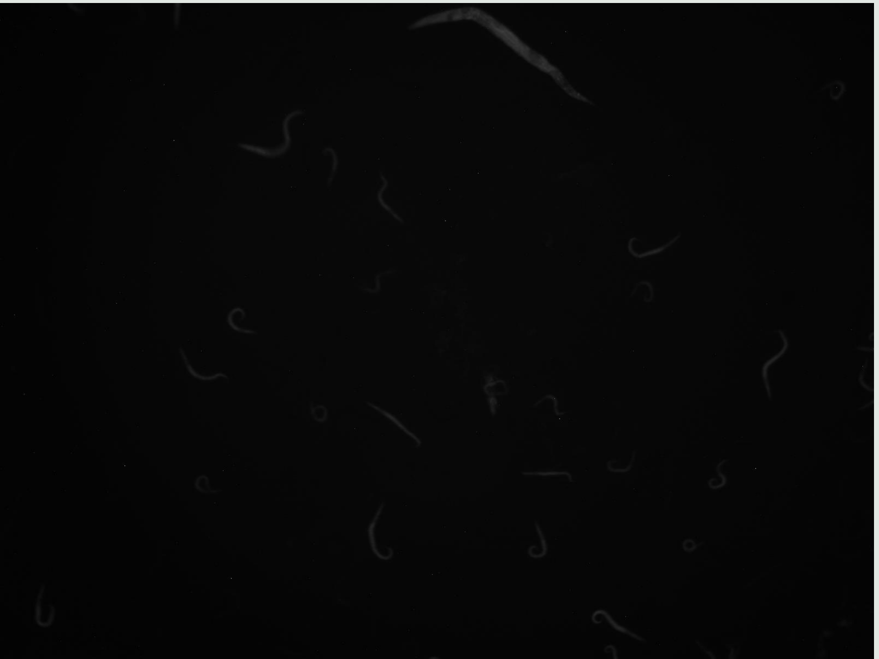

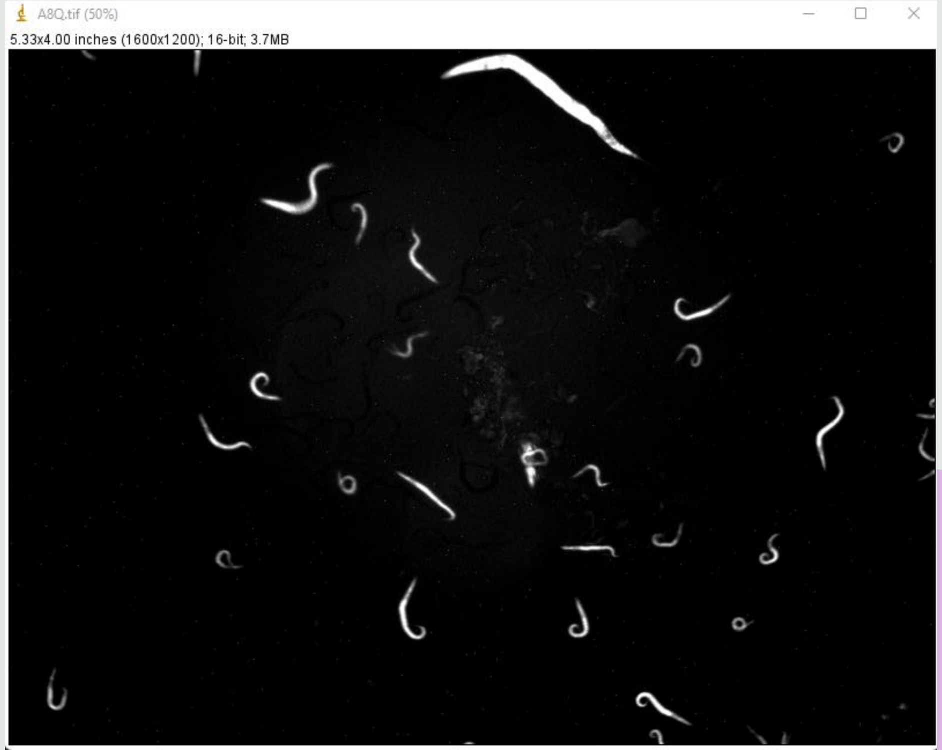

### Step 3: Use ROI selection tool to select a sample portion of the images to assay

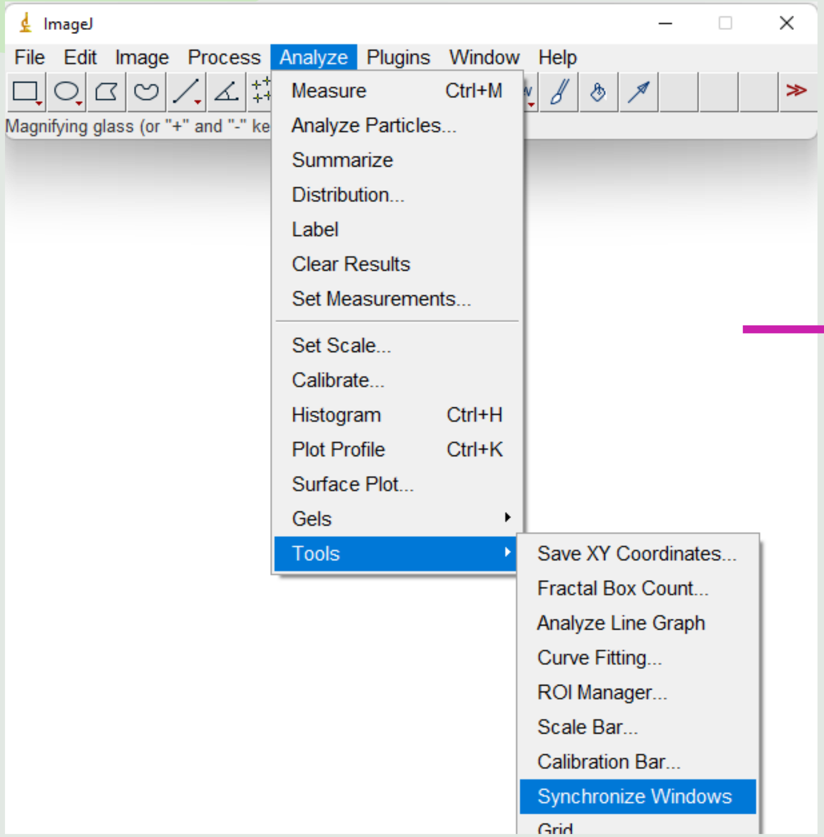


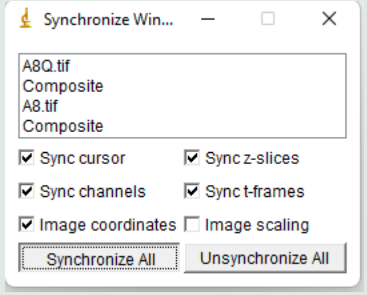


Anything you do to one image will now happen to all synchronized images

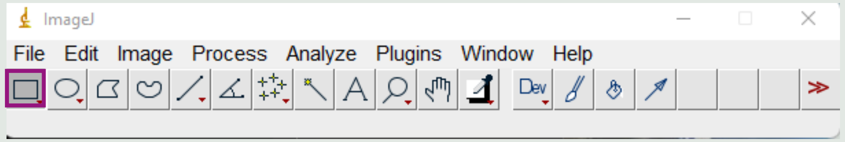


Select ROI selection tool

Then click and drag on one of the photos to select the region you want to assay. Anything selected in one photo will also be selected in the other photos

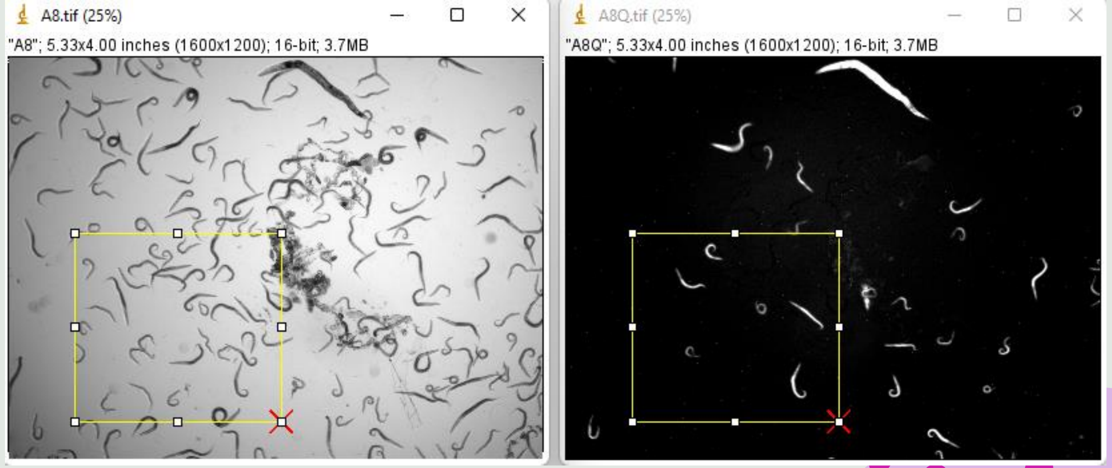


Remember to unsynchronize photos when done

You can save selection boxes and import them to other images.

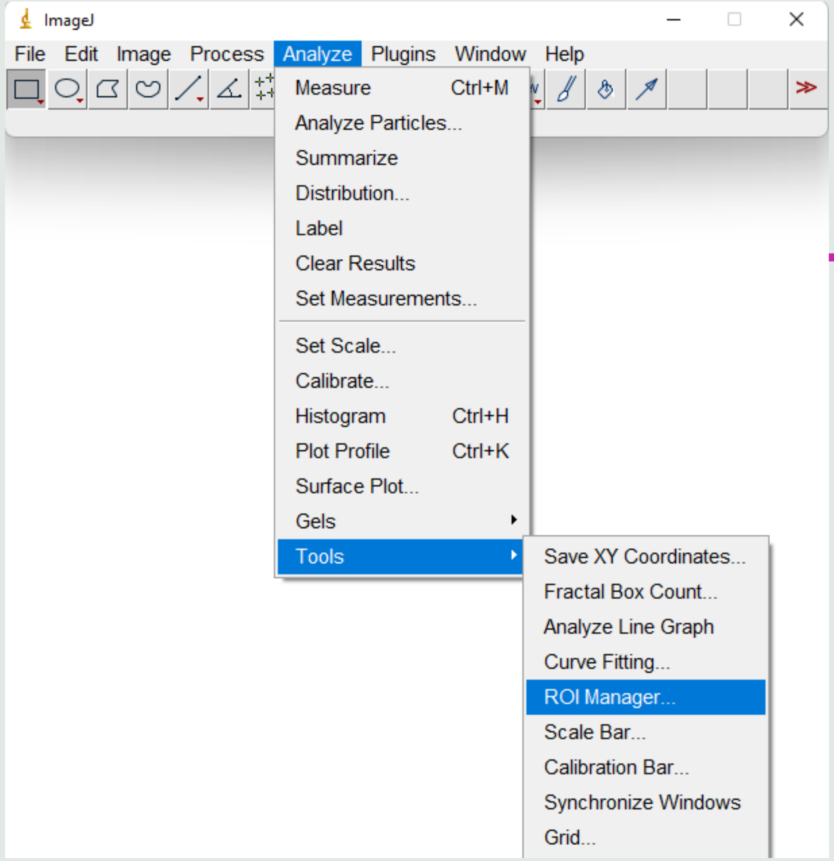


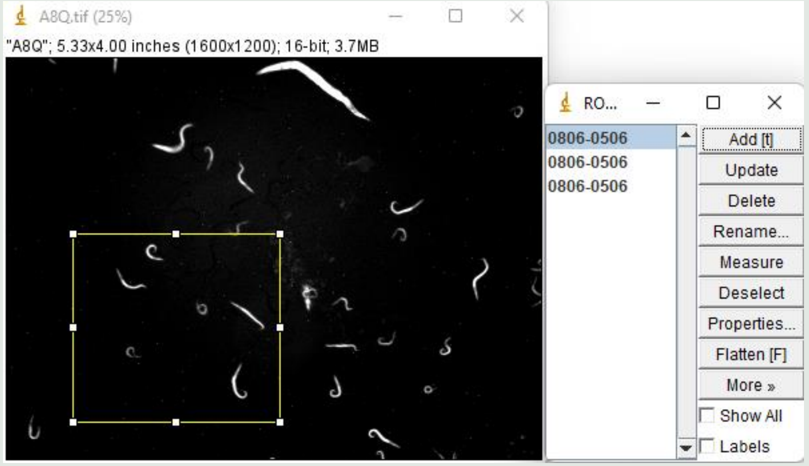


First: click on selection box and click "add" on ROI selection manager window

Second: Click "More" -> "save" to save selection box

If you restart ImageJ and want to add the saved selection box to another image: Click "More" -> "Open." If the images are the same size, the selection box will appear in the same place in every image.

### Step 4: Crop images

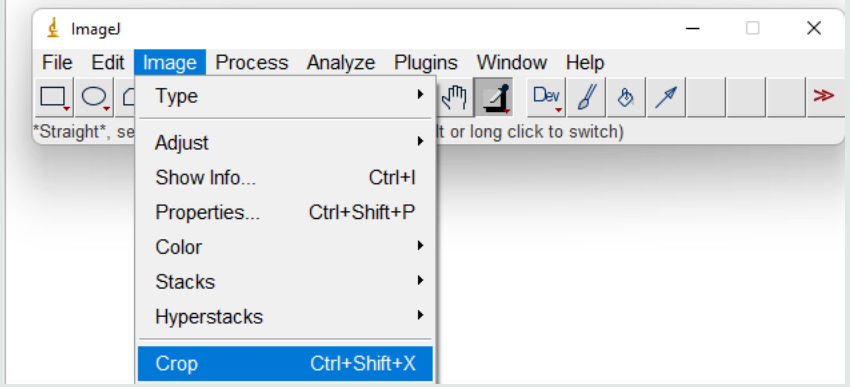


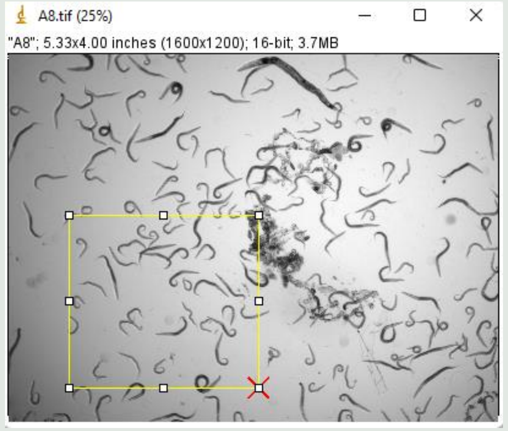


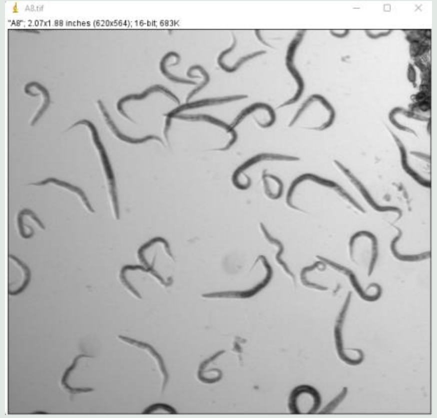


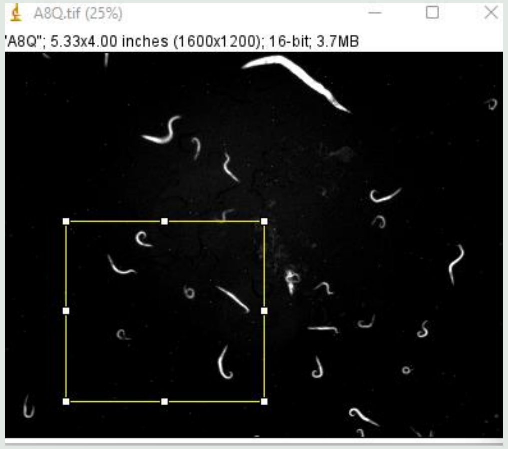


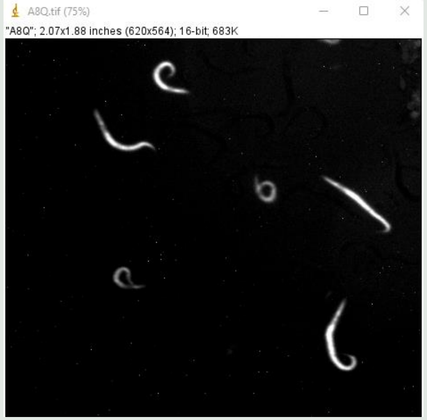

### Step 5: Colorize and merge photos

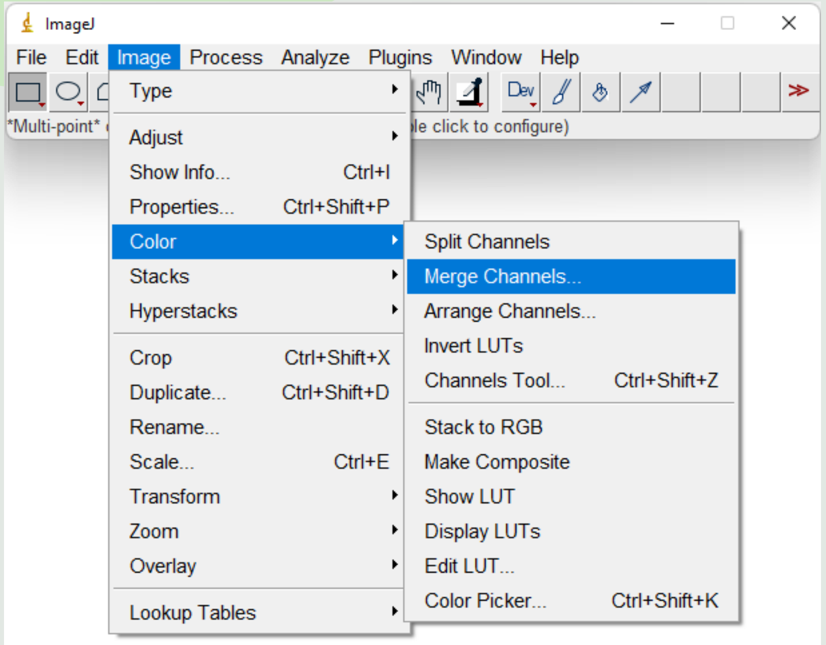


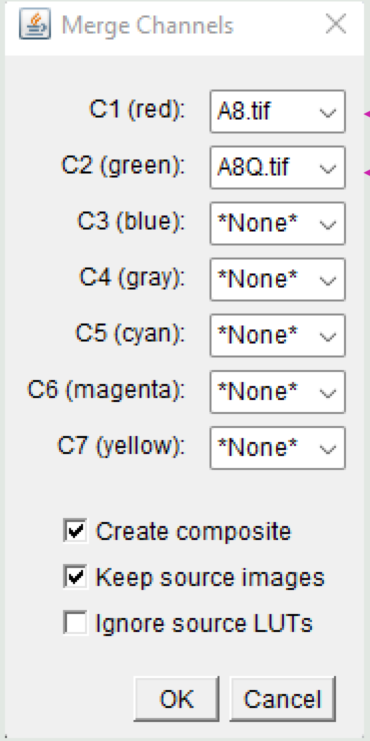


**Bright-field image set as red**

**GFP image set as green**

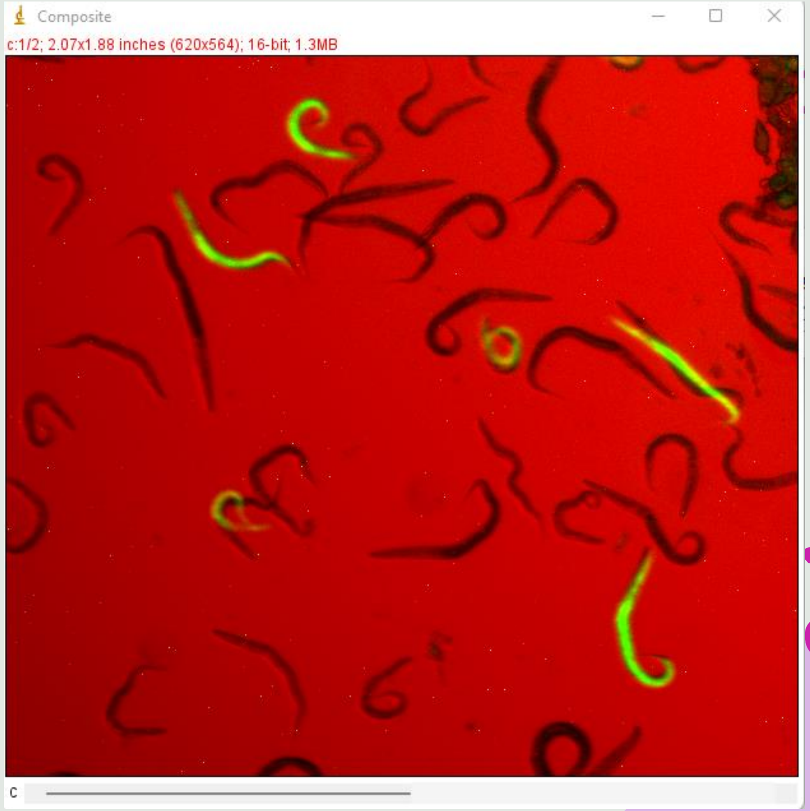


MA-line mutants are highlighted

### Now it is easy to count and compare the number of MA-line mutants to wild-type N2 worms

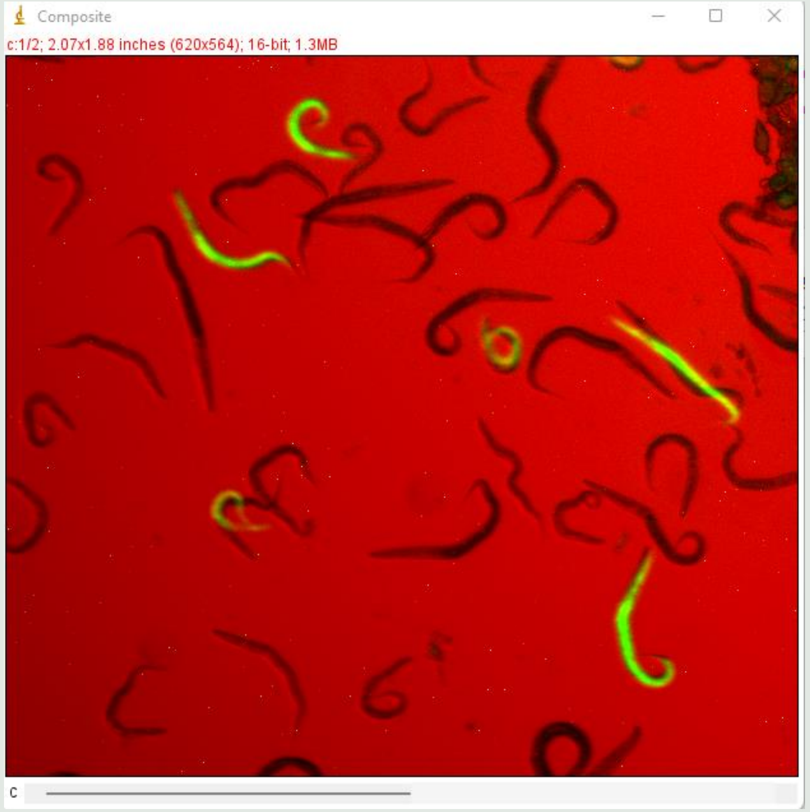


Now it is easy to count and compare the number of MA-line mutants to wild-type N2 worms

There are 7 MA-line mutants to 33 wildtype worms (not counting eggs)

The proportion found in the cropped image here should be representative of the entire population
